# Myeloid-Specific Thrombospondin-1 Deficiency Exacerbates Aortic Rupture via Broad Suppression of Extracellular Matrix Proteins

**DOI:** 10.1101/2024.07.30.605216

**Authors:** Ting Zhou, Huan Yang, Carmel Assa, Elise DeRoo, Jack Bontekoe, Brian Burkel, Suzanne Ponik, Hong S. Lu, Alan Daugherty, Bo Liu

**Author notes:** Correspondence to: Bo Liu, PhD., Ting Zhou, PhD. University of Wisconsin-Madison, 1111 Highland Avenue, WIMR 4418, Madison, WI 53705.

## Abstract

**Rationale:** Rupture of abdominal aortic aneurysms (AAA) is associated with high mortality. However, the precise molecular and cellular drivers of AAA rupture remain elusive. Our prior study showed that global and myeloid-specific deletion of matricellular protein thrombospondin-1 (TSP1) protects mice from aneurysm formation primarily by inhibiting vascular inflammation.

**Objective:** To investigate the cellular and molecular mechanisms that drive AAA rupture by testing how TSP1 deficiency in different cell populations affects the rupture event.

**Methods and Results:** We deleted TSP1 in endothelial cells and macrophages --- the major TSP1-expressing cells in aneurysmal tissues ---- by crossbreeding *Thbs1* ^flox/flox^ mice with *VE-cadherin Cre* and *Lyz2-cre* mice, respectively. Aortic aneurysm and rupture were induced by angiotensin II in mice with hypercholesterolemia. Myeloid-specific *Thbs1* knockout, but not endothelial-specific knockout, increased the rate of lethal aortic rupture by more than 2 folds. Combined analyses of single-cell RNA sequencing and histology showed a unique cellular and molecular signature of the rupture-prone aorta that was characterized by a broad suppression in inflammation and extracellular matrix production. Visium spatial transcriptomic analysis on human AAA tissues showed a correlation between low TSP1 expression and aortic dissection.

**Conclusions:** TSP1 expression by myeloid cells negatively regulates aneurysm rupture, likely through promoting the matrix repair phenotypes of vascular smooth muscle cells thereby increasing the strength of the vascular wall.

## Introduction

Aortic aneurysms are a leading cause of death in people 55 years or older, accounting for about 10,000 deaths a year in the United States^1^. Abdominal aortic aneurysm (AAA) is the most common form of aortic aneurysms; however, progressive vascular destruction and dilation may occur in the thoracic aortic segments as well. Substantial morbidity and mortality occur as the result of aneurysm rupture^2^. The complex pathophysiology of aortic aneurysm involves all layers of the aortic wall as well as the immune system. Multiple cellular processes have been identified for their essential roles in aneurysm progression, including non-resolving inflammation, excessive local production of matrix-degrading proteases, continuing destruction of structural matrix proteins, depletion and phenotypic changes of medial smooth muscle cells (SMCs), and endothelial dysfunction^3, 4, 5, 6, 7, 8^. In comparison, much less is known about the molecular and cellular mechanisms responsible for aneurysm rupture.

Thrombospondin-1 (TSP1) belongs to the thrombospondin family of matricellular proteins. TSP1 exerts a pivotal role in regulating a wide range of biological processes, including organization of extracellular matrix (ECM), facilitating cell-cell communications, and activating growth factors^9^. Multiple cell types within the vessel wall, as well as infiltrating macrophages (Mφs), are capable of expressing and secreting TSP1. In our bioinformatic analyses comparing single-cell RNA sequencing (scRNA-seq) datasets of different mouse models of aortic aneurysm and human AAA tissues, thrombospondin signaling stood out as one of the few commonly altered signaling pathways. Upregulation of the thrombospondin signaling pathway is primarily caused by increased expression of *Thbs1*, the gene encoding TSP1^10^. Furthermore, in mouse AAA models induced by local aortic injuries with elastase or CaCl_2_, global or myeloid deletion of *Thbs1* protected mice from developing aneurysm^11, 12^. Mechanistic studies indicated the absence of aneurysm formation in *Thbs1* deficient mice resulted from diminished inflammation due to insufficient Mφ migration^11, 12^. However, whether high abundance of aortic TSP1 exerts any role in aneurysm rupture remains unknown because neither elastase nor CaCl_2_ models produce aortic rupture^4, 13^. In the current study, we tested the hypothesis that TSP1 depletion from different cellular sources may lead to distinct effects on aneurysm rupture. To create an aneurysm with aortic rupture, we induced two comorbidities in mice --- hypercholesterolemia and hypertension --- by combining feeding a Western diet with infection with of AAVs encoding a mouse PCSK9 gain-of-function mutation (PCSK9D377Y)^14^ and angiotensin II (AngII) infusion through osmotic pumps^15^. Within a period of 28 days, myeloid-specific *Thbs1* deficient mice (*Thbs1^Δ^*^Mφ^) exhibited aortic rupture at a rate that was 2.6-fold higher than *Thbs1*^wt^. Intriguingly, *Thbs1^Δ^*^Mφ^ mice that survived AngII infusion showed smaller aneurysmal dilation than *Thbs1*^wt^. Endothelial-specific *Thbs1* deficient mice (*Thbs1^Δ^*^EC^) had a moderate protective effect on aneurysm formation but did not alter the rate of rupture. Mechanistically, scRNA-seq analyses revealed unique cellular and molecular signatures of *Thbs1^Δ^*^Mφ^ aortas characterized by suppressed inflammation and diminished extracellular matrix production accompanied by reduction of fibroblasts as well as a phenotypic shift of smooth muscle cells. Spatial transcriptomics on human AAA tissue further confirmed lower TSP1 abundance at aortic dissection sites.

## Methods

### Data Availability

All raw data and analytical methods are available from the corresponding author on reasonable request. Sequencing data will be made publicly available at the Gene Expression Omnibus upon publication. The computation code used in this study is available on request.

### Human AAA tissues

Human aortic aneurysm tissues were obtained from patients undergoing either elective open AAA repair (n=4), or emergency open ruptured AAA repair (n=4). The protocol for collecting human tissue samples was approved by the Institutional Review Board at the University of Wisconsin – Madison (ID: 2021-0215) as a minimal risk IRB, thus patient demographic information was not permitted to be collected. All experiments conducted with human tissue samples were performed in accordance with the relevant guidelines and regulations.

### Mice

All animal studies were performed with the approval of the Institute Animal Care and Use Committee at the University of Wisconsin – Madison (Protocol #M005792). *Thbs1^-/-^* (RRID: IMSR_JAX:006141) and *Apoe^-/-^* mice (RRID: IMSR_JAX:002052) were obtained from The Jackson Laboratory and crossbred to generate *Thbs1^-/-^Apoe^-/-^* and *Thbs1^+/+^Apoe^-/-^* mice. Endothelial *Thbs1* deficient mice (*Thbs1^fl/fl^/VE-CRE*) were generated by crossing *Thbs1* flox mice^12^ with *VE-cadherin Cre* mice (RRID: IMSR_JAX:006137), *Thbs1^wt/wt^/VE-CRE* were used as control. Myeloid *Thbs1* deficient mice (*Thbs1^fl/fl^/Lyz2-Cre*) and *Thbs1^wt/wt^/Lyz2-Cre* control were generated previously^12^ by crossing *Thbs1* flox mice with *Lyz2-Cre* mice (RRID: IMSR_JAX:004781). Both sexes were used; however, most experiments were performed on male mice due to the lack of aneurysm development in female mice. All mice were maintained on standard laboratory diet (2018 Teklad global 18% protein rodent diets, ENVIGO) unless otherwise indicated.

### PCSK9 AAV AngII model of AAA

Mice at 8-12 weeks of age were intraperitoneally injected with 20×10^10^ genome copies of adeno-associated virus (AAV) expressing the mouse PCSK9 gain-of-function mutation D377Y, as described previously^14^. Two weeks later, mice with serum cholesterol concentrations exceeding 250 mg/dL were subcutaneously implanted with Alzet osmotic minipumps (Model 2004, DURECT Corporation) infusing AngII (1000 ng/kg/min, Sigma-Aldrich A9525) or PBS vehicle. The mice were fed with a Western diet (TD.88137, ENVIGO) starting from the time of AAV injection and throughout the study. An aneurysm was defined as an increase of 50% or greater in the external width of the suprarenal aorta, as compared with that of the infrarenal region. The abdominal aortas were either snap frozen in liquid nitrogen, embedded in optimal cutting temperature compound (Sakura Tissue Tek) and cut into six-micrometer cross sections, or subjected to single-cell dissociation followed by RNA sequencing.

### Modified CaCl_2_ Model

Twelve to 16 weeks old male mice were anesthetized by isoflurane inhalation. Infrarenal abdominal aorta was isolated following a midline incision. A small piece of gauze soaked in CaCl_2_ (0.5 mol/L) was applied perivascularly for 10 minutes. The gauze was replaced with another piece of PBS-soaked gauze for 5 minutes. Mice in sham group were immersed in NaCl (0.5 mol/L) for 10 minutes followed by PBS-soaked gauze for 5 minutes. After 28 days, mice were euthanized and perfused with PBS, and the maximum external diameter of the infrarenal aorta was measured using a digital caliper. In this model, AAA forms at the infrarenal segment of the abdominal aorta. An aneurysm was defined as a 50% increase in the maximum external aortic diameter as compared with that before CaCl_2_ or NaCl exposure.

### Blood pressure measurements

Blood pressure was measured on conscious mice using a non-invasive tail-cuff system (CODA Monitor, Kent Scientific Corporation) as described previously^16^. Data were collected and analyzed based on 20 measurements of each mouse. Mean blood pressure of each mouse was used for data analysis.

### Cholesterol assay

Blood samples were collected using retro-orbital bleeding on mice sedated with isoflurane. Serum cholesterol concentrations were determined using an enzymatic kit (Cat # 439-17501; Wako Chemicals USA).

### Ultrasound imaging and analysis

Aneurysm growth was monitored by transabdominal ultrasound scanning (40 MHz, Vevo 2100, VisualSonics) prior to (baseline) and post AAA induction. Internal and external aortic diameters were measured and analyzed. Circumferential strain was calculated as [(D_s_/D_d_)^2^-1]/2, where D_s_ was systolic internal diameter, and D_d_ was diastolic internal diameter^17^.

### RNA *in situ* hybridization

Fresh frozen sections of murine aortas were analyzed using the Advanced Cell Diagnostics RNAscope Fluorescent Multiplex Reagent Kit (#320850) with RNAscope probes against mouse *Thbs1* (457891-C3) and *Pecam1* (316721). Tissue incubations, probe hybridization, and signal amplification steps were conducted following the manufacturer’s instructions. Images were captured using a Nikon A1RS confocal microscope system, and subsequent analysis was performed using Fiji/ImageJ software.

### Immunofluorescent staining

PFA-fixed tissue sections were blocked with normal donkey serum (10% vol/vol) and incubated at 4 °C overnight with primary antibodies anti-F4/80 (123102, BioLegend), anti-Col1a1 (NB600-408, Novus Biologicals), anti-MYH11 (ab53219, abcam), anti-TSP1 (AF3074, R&D Systems). Normal goat or rabbit IgG was used as negative controls. After three washes with PBS, sections were incubated with Alexa Fluor 488- or Alexa Fluor 594-conjugated secondary antibodies for 1 hour at room temperature. DAPI was used to stain nuclei. Images were acquired with a Nikon A1RS confocal microscope system and analyzed using ImageJ software. Tortuosity index was measured by NeuronJ plugin, Tortuosity index=True trace of elastin fiber/Euclidian trace of elastin fiber.

### Mφ-SMC co-culture

Bone marrow cells were collected from femurs and tibias of mice and differentiated into bone marrow–derived Mφs (BMDMs) using Mφ medium (DMEM with FBS (10% vol/vol), penicillin/streptomycin (1% wt/vol), and mouse M-CSF (20 ng/mL)). Medium was changed after 3 days of incubation at 37 °C, 5% CO_2_. After a further 4 days of culture, the adherent cells were stimulated with LPS (100 ng/mL) and IFNγ (20 ng/mL) for 4 hours, then seeded in the upper compartment of Transwell (3450, Corning). Mouse aortic smooth muscle cell line (MOVAS, CRL-2797, ATCC) were seeded in the lower compartment. After 24 hours of co-culture, MOVAS was collected and *Thbs1* mRNA was analyzed.

### Single-Cell RNA Sequencing and Analysis

Seven days into AngII infusion, mouse infrarenal abdominal aortas of *Thbs1*^wt^ and *Thbs1*^ΔMφ^ (n=5 for each group) were collected and digested sequentially in two digestion buffers (PBS containing collagenase I (200U/mL; SCR103, Sigma Aldrich), elastase 0.05 U/mL; (E1250, Sigma Aldrich), neutral protease (5 U/mL, LS02111, Worthington), and deoxyribonuclease I (0.3 U/mL, M6101, Promega)^18^ for 20 minutes, followed by DMEM containing collagenase type II (5 mg/mL, C6885, Sigma-Aldrich) and elastase (0.5 mg/mL, LS002292, Worthington Biochemistry]) for 2.5 minutes at 37°C)^19^. The tissue suspension was filtered with a 40 μm cell strainer, then centrifuged at 500 g for 5 minutes. Cells were resuspended with PBS containing BSA (0.04% wt/vol). Single-cell suspensions from 5 *Thbs1*^wt^ or 5 *Thbs1*^ΔMφ^ infrarenal abdominal aortas were pooled together as one sample. Ten thousand cells per sample were loaded on a Chromium Controller (10x Genomics). The scRNA-seq libraries were constructed using the Chromium Single Cell 3’ v3.1 Reagent Kit according to the manufacturer’s guidelines (10x Genomics). cDNA libraries were uniquely sample indexed and pooled for sequencing. A MiSeq (Illumina) sequencing run was used to sample balance on a NovaSeq S2 flowcell (Illumina) using a 2×50 bp sequencing reaction targeting >90,000 reads/cell.

Raw Illumina sequencing reads were aligned to mm10 (GENCODE vM23/Ensembl 98) reference genome and, subsequently, genes were quantified as UMI counts using Simpleaf (alevin-fry pipline, V0.9.0)^20^. Downstream analysis including quality control, sample integration, clustering, differential expression and pathway enrichment analysis was performed by scvi-tools V1.0.4^21^ and Scanpy V1.1.0^22^ on python 3.11.9 platform. RNA velocity analysis was performed by scVelo V0.3.1^23^. Cell–cell communication analysis was performed using CellChat 2.1.2^24^ on R 4.4.0.

### Spatial transcriptomics and analysis

Spatial transcriptomics of human AAA tissues (4 elective AAA, 4 ruptured AAA) were conducted using Visium Spatial for FFPE Gene Expression Kit (1000338, 10x Genomics) according to the manufacturer’s guidelines. Raw sequencing fastq files were processed by using Space Ranger 2.0.0 standard pipeline and further analyzed using Squidpy v1.3.1^25^ and chrysalis v0.2.0 (bioRxiv, doi:10.1101/2023.08.17.553606).

### Statistical Analyses

ScRNA-seq analysis was described above. Other results were presented as mean±SD. Continuous variables were assessed for normality using the Shapiro-Wilk normality test. Data not exhibiting a normal distribution were log2-transformed and retested for normality. Equality of variance was examined by an F test. Two-tailed Student’s t test for normally distributed and equivalent variance data and Mann-Whitney nonparametric test for skewed data that remained deviated from normality after transformation were used to compare between 2 conditions. Statistical analyses were performed using GraphPad Prism 10 (GraphPad Software). Experiments were repeated as indicated. Differences with *P*<0.05 were considered statistically significant.

## Results

### Global TSP1 deficient mice in the ApoE null background responded to AngII with an increased rate of aneurysm rupture

To investigate the role of TSP1 in aortic aneurysm under hypercholesterolemia and hypertension conditions --- the comorbidities found in most AAA patients, we generated TSP1 and apolipoprotein E (ApoE) compound deficient mice by breeding *Thbs1*^-/-^ with *Apoe*^-/-^ mice. We infused male mice with AngII (1000 ng/kg/min) via osmatic pumps, and collected aortas after 28 days. Two out of 13 (15.4%) *Thbs1*^+/+^/*Apoe*^-/-^ mice died from aortic rupture before the end of experiment, while 6 out of 11 (54.5%) *Thbs1*^-/-^/*Apoe*^-/-^ mice succumbed to aortic rupture (Figure S1A). Among the mice that survived 28 days AngII infusion, there was no significant difference in the aortic diameter between *Thbs1*^+/+^/*Apoe*^-/-^ and *Thbs1*^-/-^/*Apoe*^-/-^ (Figure S1B-D).

### Mφ-, but not EC-, specific TSP1 deficiency affected aortic rupture

Several cell types, including endothelial cells (EC) and Mφs, within the aneurysmal aortic wall express TSP1^10,26^. We generated cell-specific TSP1 deficient mice (*Thbs1*^ΔEC^) by crossing *Thbs1^flox/flox^* mice with *VE-cadherin Cre*. *In situ* hybridization validated the successful deletion of *Thbs1* specifically in endothelial cells (Figure S2). Due to the lack of Mφ-specific *Cre*, myeloid-specific TSP1 deficient mice (*Thbs1^flox/flox^/Lyz2 Cre*) were used as a surrogate for Mφ-specific *Thbs1* deficient (*Thbs1*^ΔMφ^). Efficiency and specificity deletion of *Thbs1*^ΔMφ^ have been reported in our previous publication^12^.

Both *Thbs1*^ΔEC^ and *Thbs1*^ΔMφ^, males and females, along with their corresponding TSP1 wildtype mice, were subjected to hypercholesterolemia induction by infection of AAV.PCSK9D377Y (20×10^10^ genome copies per mouse) and Western diet followed by AngII (1,000 ng/kg/min) infusion (Figure 1A and 2A).

**Figure 1.**
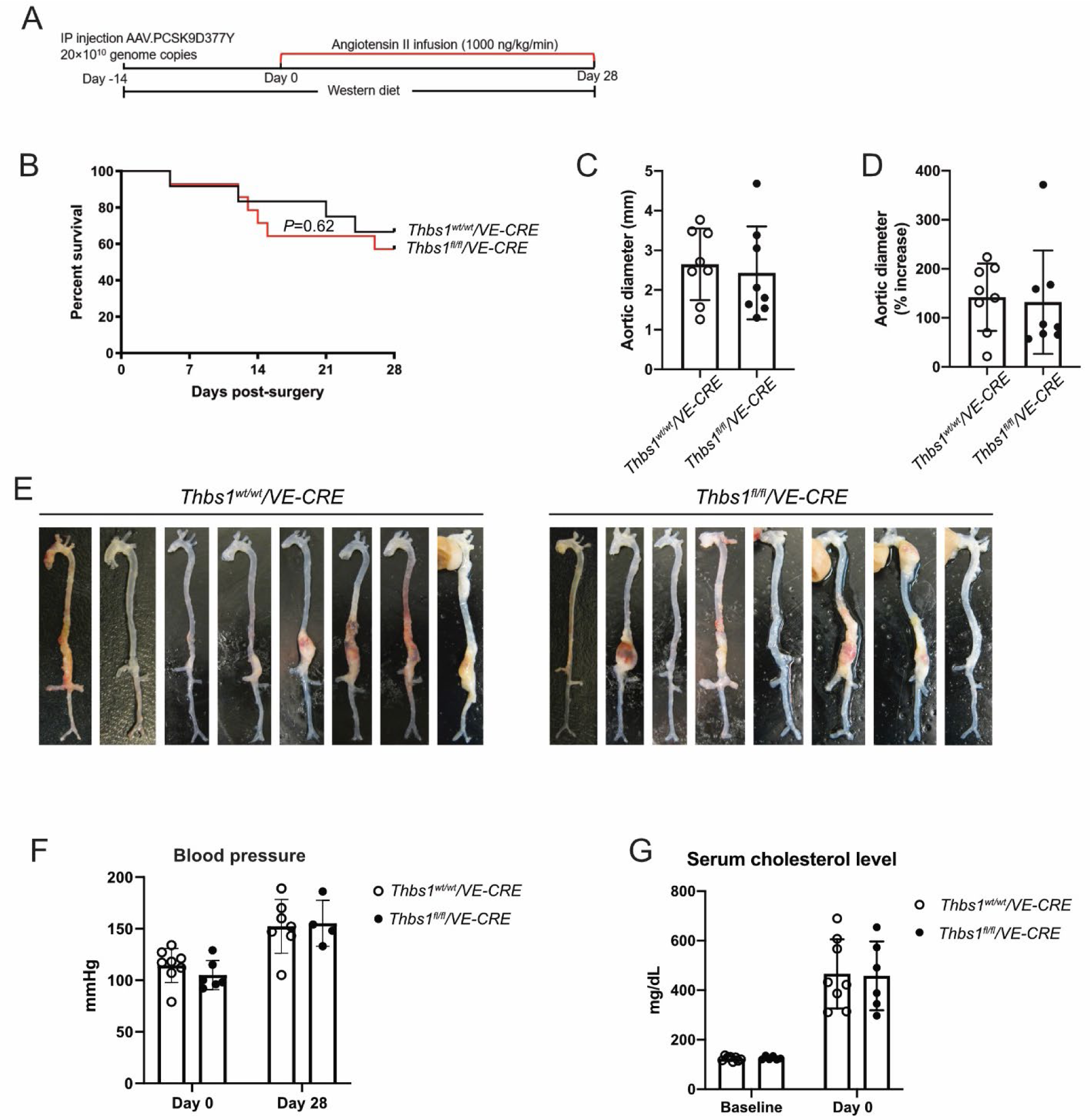
Endothelial-specific *Thbs1* deficiency did not augment aortic rupture. **(A)** Scheme of the AAV.PCSK9D377Y-angiotensin II (Ang II) model. **(B)** Survival curve of *Thbs1^wt/wt^/VE-CRE* and *Thbs1^fl/fl^/VE-CRE* mice during 28 days of Ang II infusion (1000 ng/kg/min). **(C&D)** Maximum external aortic diameter at the suprarenal abdominal aorta in survived mice (C), and the percentage increase compared to the maximum external aortic diameter of the infrarenal region (D). **(E)** Images of the aortas taken from mice survived the full course of Ang II infusion. **(F)** Blood pressure measured using the tail-cuff system before Ang II infusion (Day 0) and on Day 28 of Ang II infusion. **(G)** Serum cholesterol levels before AAV injection (baseline) and 14 days after AAV injection (Day 0 of Ang II infusion). N=12 for *Thbs1^wt/wt^/VE-CRE* and n=14 for *Thbs1^fl/fl^/VE-CRE* were subjected to Ang II infusion. Log-rank test was performed in (B). Data in (C), (D), (F), (G) were presented as mean±SD. Two-tailed Student t test was performed in (C) and (D). Two-way ANOVA followed by Tukey’s multiple comparisons was performed in (F) and (G).

**Figure 2.**
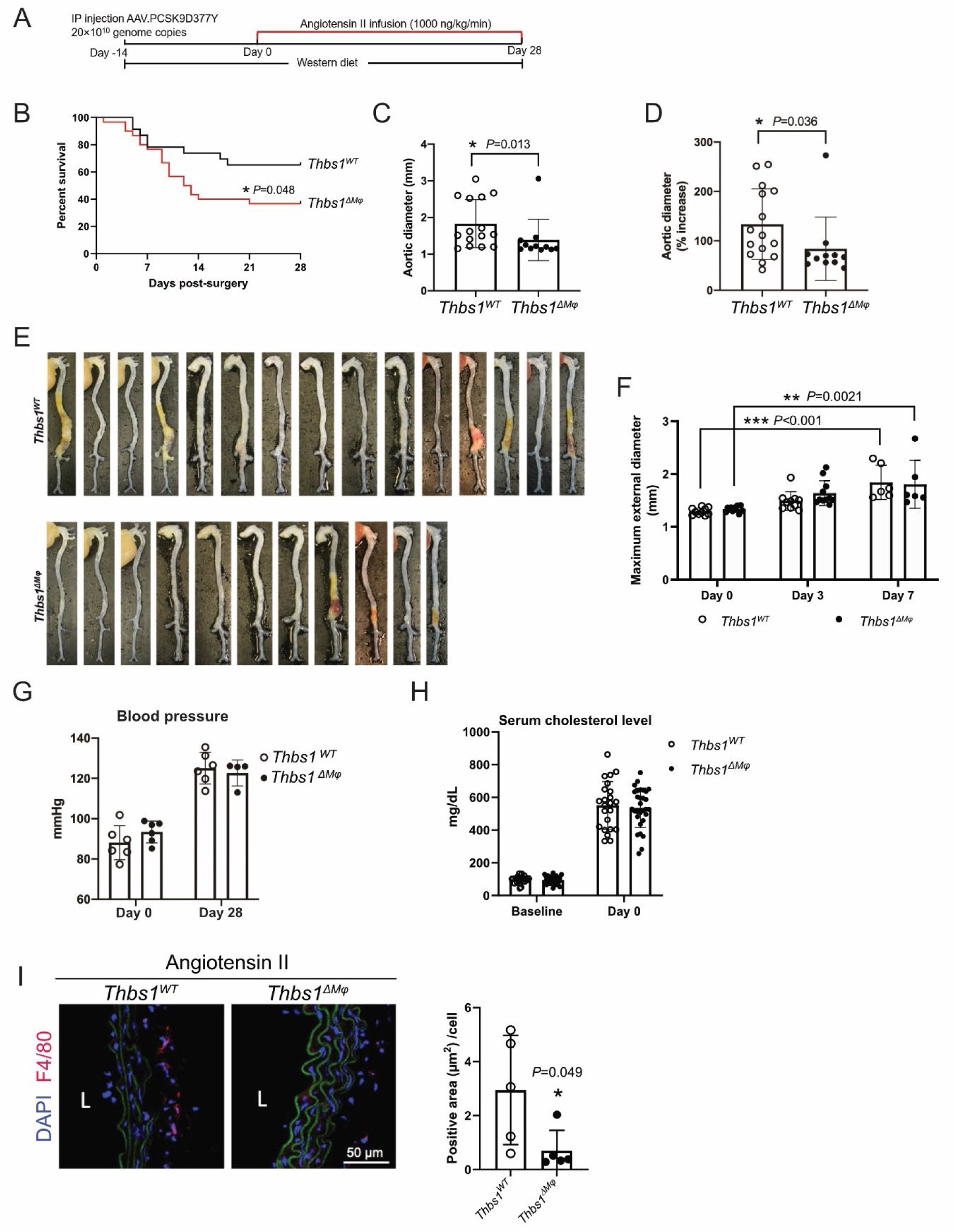
Myeloid-specific *Thbs1* deficiency augmented aortic rupture. **(A)** Scheme of the AAV.PCSK9D377Y-angiotensin II (Ang II) model. **(B)** Survival curve of *Thbs1^wt/wt^/Lyz2 Cre* (*Thbs1^WT^*) and *Thbs1^flox/flox^/Lyz2 Cre* (*Thbs1^ΔMφ^*) mice during 28 days of Ang II infusion (1000 ng/kg/min). **(C&D)** Maximum external aortic diameter at the suprarenal abdominal aorta in survived mice (C), and the percentage increase compared to the maximum external aortic diameter of the infrarenal region (D). **(E)** Images of the aortas taken from mice survived the full course of Ang II infusion. **(F)** Maximum external aortic diameter at the suprarenal abdominal aorta traced by ultrasound scanning. **(G)** Blood pressure measured using the tail-cuff system before Ang II infusion (Day 0) and on Day 28 of Ang II infusion. **(H)** Serum cholesterol levels before AAV injection (baseline) and 14 days after AAV injection (Day 0 of Ang II infusion). **(I)** Immunostaining and quantification of aortic cross-sections 3 days into Ang II infusion using anti-F4/80. Green: elastin autofluorescence. “L” indicates the aortic lumen. A total of 23 *Thbs1^WT^* and 30 *Thbs1^ΔMφ^* were subjected to Ang II infusion. Log-rank test was performed in (B). Data in (C-D) and (F-I) were presented as mean±SD. Two-tailed Student t test was performed in (C), (D), and (I). Two-way ANOVA followed by Tukey’s multiple comparisons was performed in (F-H). \**P*<0.05, \*\**P*<0.01, \*\*\**P*<0.001.

Noticeably, only ∼50% of female mice responded to AAV-PCSK9 and Western diet feeding with elevated blood cholesterol > 250 mg/dL, in contrast to ∼95% in male mice. Even among the female mice that developed hypercholesterolemia, the increased concentrations were lower than that noted in male mice. All female mice survived the 4-week AngII infusion. Measurements of aortic diameter during euthanization on day 28 showed that none of the female mice, regardless of genotype, developed aortic aneurysm (n=6 for *Thbs1*^wt^, n=8 for *Thbs1*^ΔMφ^) (Figure S3). Due to the lack of aneurysm response in females, the rest of the studies were only performed in males.

Most male TSP1 wildtype mice developed aneurysm, albeit with different severity. Specifically, 6 out of 14 *Thbs1*^ΔEC^ and 4 out of 12 wildtype controls (*Thbs1^wt/wt^/VE-cadherin Cre*) died from aortic rupture before the last day of AngII infusion; however, the rate of rupture was not significantly different between the genotypes (Figure 1B). Among the mice who survived to day 28 of AngII infusion, there was a trend of reduction in aneurysm size in the *Thbs1*^ΔEC^ group; however, the difference in aortic dilation did not reach statistical significance due to a single unusually large aneurysm in the deficient group (maximum external aortic diameter *Thbs1*^ΔEC^ vs wildtype: 132±99% vs 142±64%) (Figure 1C-E). Furthermore, the loss of TSP1 in ECs did not alter blood pressure or cholesterol concentrations (Figure 1F&G). The potential contribution of endothelial TSP1 to aneurysm pathophysiology was also demonstrated in the CaCl_2_-induced aneurysm model, a model that does not involve hypercholesterolemia or hypertension. Compared to the wildtype littermates, *Thbs1*^ΔEC^ developed significantly less aortic dilation (external diameter % increase 28 days after AAA induction: *Thbs1*^ΔEC^ vs wildtype = 18.45±9.46% vs 47.34±14.74%) and lower arterial stiffness (circumferential strain 28 days after AAA induction: *Thbs1*^ΔEC^ vs wildtype = 0.141±0.039 vs 0.077±0.028, Figure S4).

We next examined the effect of myeloid-specific *Thbs1* deletion on the AngII response and observed that a significantly higher number of *Thbs1*^ΔMφ^ males died from aortic rupture before scheduled euthanization on day 28 of AngII infusion than their wildtype control (*Thbs1^wt/wt^/Lyz2 Cre*): 19 out of 30 vs 8 out of 23 (Figure 2B). Paradoxically, survived *Thbs1*^ΔMφ^ mice showed a significantly smaller aortic expansion than wildtype (*Thbs1*^ΔMφ^ vs wildtype: 84±61% vs 133±69%) (Figure 2C-E). Ultrasound imaging, performed before AngII infusion (baseline), and after 3 and 7 days of AngII infusion showed that, 7 days of AngII infusion induced significant aortic dilation in both *Thbs1*^ΔMφ^ and wildtype mice, but there was no difference in aortic diameter between genotypes (Figure 2F). Like the EC-specific deficiency, the loss of TSP1 in myeloid cells did not alter blood pressure or cholesterol concentrations (Figure 2G&H)

### Cellular characteristics of the rupture-prone aortic tissues

Given aneurysm rupture is the most dangerous outcome of AAAs, we focused the rest of investigation on the molecular and cellular mechanism by which myeloid-specific *Thbs1* deficiency exacerbates aortic rupture triggered by AngII. Because our prior work implicated TSP1 in Mφ mobility^12^, we next examined Mφ accumulation by immunostaining and found that the aortic wall of *Thbs1*^ΔMφ^ mice had significantly fewer Mφs (F4/80+) compared to wildtype mice (Figure 2I).

To further investigate how lacking myeloid TSP1 affects the aortic wall, we conducted scRNA-seq on abdominal aortas collected after 7 days of AngII infusion, a time interval before most cases of rupture occurred. A total of 5624 cells from the *Thbs1*^WT^ group and 5401 cells from the *Thbs1*^ΔMφ^ group, each pooled from 5 aortas, were recovered after sequencing. Unsupervised clustering revealed the presence of multiple cell populations in the aortic wall (Figure 3A), which were identified---based on the expression of canonical marker genes --- as SMCs, fibroblasts, myofibroblasts, ECs, Mφs, dendritic cells, and T/NK/B cells (Figure 3A-C). Consistent with the immunostaining (Figure. 2I), scRNA-seq analyses showed *Thbs1*^ΔMφ^ contained fewer Mφs than wildtype aortas (*Thbs1*^ΔMφ^ vs wildtype= 5.7% vs 15.1%) (Figure 3D). Additionally, *Thbs1*^ΔMφ^ increased the percentage of SMCs (*Thbs1*^ΔMφ^ vs wildtype= 73.5% vs 49.0%) but decreased the percentage of fibroblasts (*Thbs1*^ΔMφ^ vs wildtype= 12.0% vs 24.3%) (Figure 3D).

**Figure 3.**
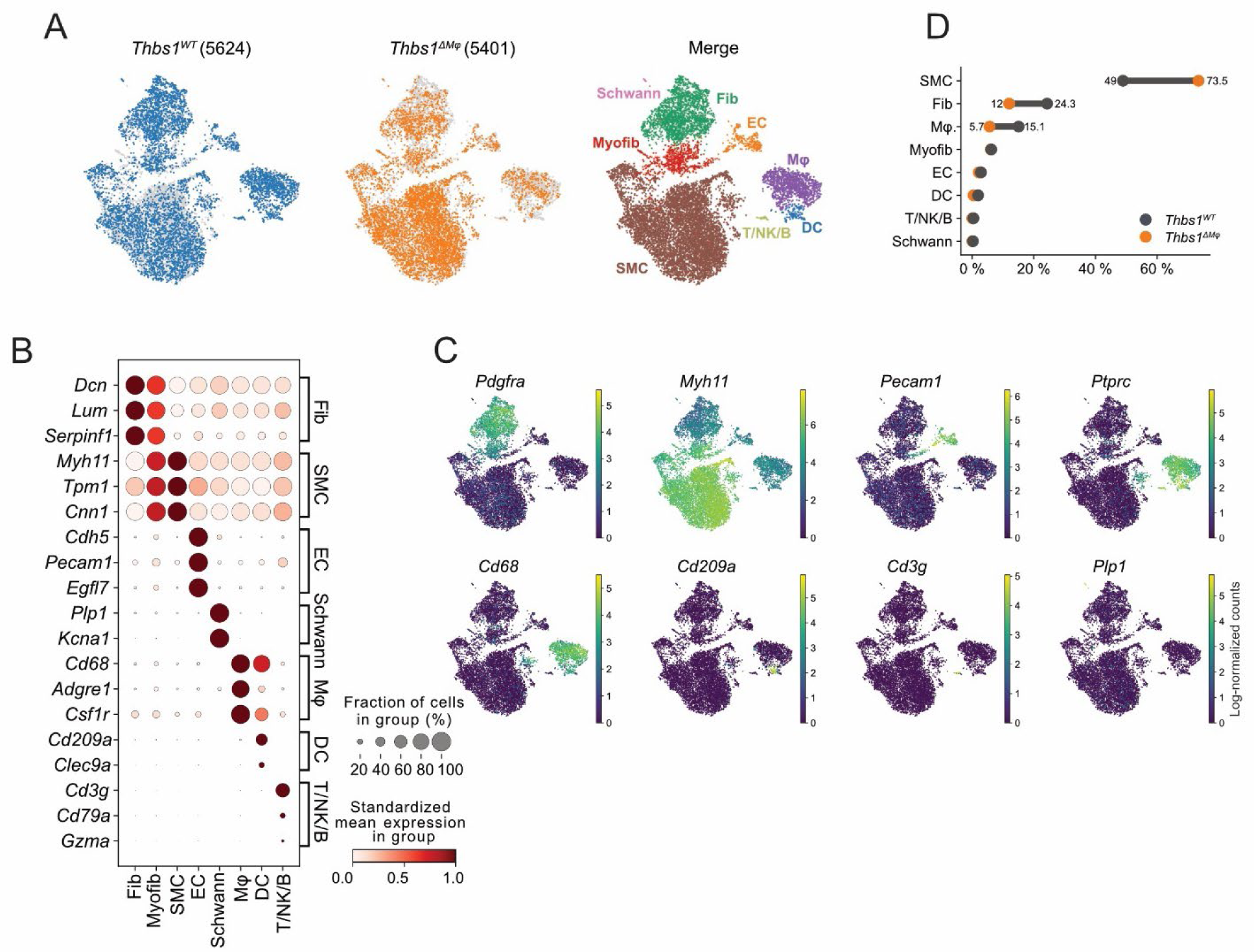
Single-cell RNA sequencing revealed the cellular characteristics of the rupture-prone aortic tissues. After 7 days of angiotensin II infusion, infrarenal abdominal aortas of *Thbs1^WT^* and *Thbs1^ΔMφ^* (n=5 for each group) were collected and analyzed by single-cell RNA sequencing. **(A)** Uniform manifold approximation and projection (UMAP) plots of cell populations presented in *Thbs1^WT^* and *Thbs1^ΔMφ^* and combined (merge). **(B)** Expression of canonical cell markers in each cell type. **(C)** UMAP plots of cell markers in *Thbs1^WT^* and *Thbs1^ΔMφ^* data combined. **(D)** Percentages of main cell types in *Thbs1^WT^* and *Thbs1^ΔMφ^*.

### Transcriptomic profile of the rupture-prone aortic tissues

Differentially expressed gene (DEG) analysis showed that myeloid TSP1 deficiency increased abundance of 402 genes, but decreased the abundance of 537 genes (Figure 4A). Gene set enrichment analysis suggests that upregulated genes clustered mostly in pathways responsible for smooth muscle contraction and the downregulated genes in pathways responsible for extracellular matrix (ECM) synthesis/organization and inflammation (Figure 4B&C). The type I collagen coding gene (*Col1a1*) was among the most downregulated genes in *Thbs1*^ΔMφ^ (Figure 4A&C).

**Figure 4.**
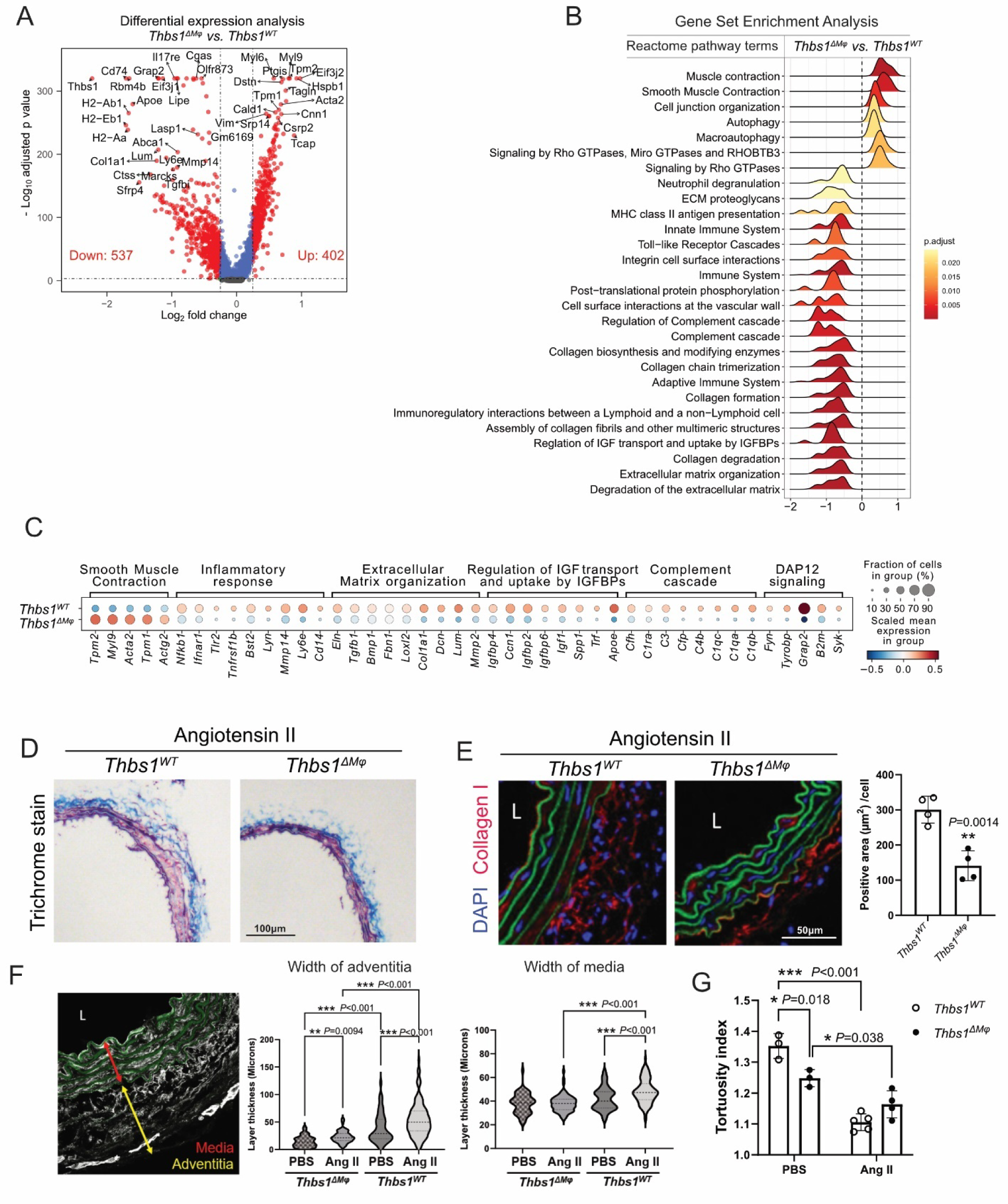
Transcriptomic profile of the rupture-prone aortic tissues. **(A)** Volcano plot of differentially expressed genes (DEGs) in *Thbs1^ΔMφ^* versus *Thbs1^WT^* group. **(B)** Gene Set Enrichment Analysis (GSEA) of up- and down-regulated gene sets in *Thbs1^ΔMφ^* versus *Thbs1^WT^* group. **(C)** Gene expression in the top altered gene set and signaling pathways in *Thbs1^WT^* and *Thbs1^ΔMφ^* aortic tissues. **(D)** Trichrome staining of aortic cross-sections in *Thbs1^WT^* and *Thbs1^ΔMφ^* mice 7 days into angiotensin II (Ang II) infusion. **(E)** Immunostaining and quantification of aortic cross-sections after 7 days of Ang II infusion using anti-Col1a1. Green: elastin autofluorescence. “L” indicates the aortic lumen. **(F)** Width of media and adventitia was measured using images from (E). Green: elastin autofluorescence. **(G)** Elastin curvature (tortuosity index) was measured using images from (E). Data were presented as mean±SD. Two-way ANOVA followed by Tukey’s multiple comparisons was performed. Two-tailed Student t test was performed in (E). One-way and two-way ANOVA followed by Tukey’s multiple comparisons were performed in (F) and (G), respectively. \**P*<0.05, \*\**P*<0.01, \*\*\**P*<0.001.

Both trichrome staining and immunostaining for Col1a1 confirmed the reduction of collagen in *Thbs1*^ΔMφ^ aortic tissues of AngII-infused mice (Figure 4D&E). Compared to wildtype aortic wall, *Thbs1*^ΔMφ^ aorta exhibited much thinner adventitia in both PBS control and AngII-infused mice (Figure 4F). The thickness of medial layer was also reduced by the conditional knockout, but only in the AngII-infused mice (Figure 4F). Furthermore, the curviness of elastin fibers (tortuosity), a measure of elastin deposition and assembly, was significantly decreased in *Thbs1*^ΔMφ^ mice incubated with PBS (Figure 4G). AngII stimulation profoundly reduced the curviness of elastin bundles in the mice regardless of their genotypes (Figure 4G).

UMAP plots showed that downregulated expression of *Thbs1* in multiple cell types including Mφs and SMCs of mutant aortas (Figure S5A). However, *Lyz2* was mainly expressed by Mφs (Figure S5B), which is consistent with the literature^27^. Because TSP1 is a major signal sent by inflammatory Mφs to SMCs^10^, we hypothesize that Mφs, with or without TSP1, have different effects on SMCs, including the expression of *Thbs1*. To evaluate this hypothesis, we isolated Mφs from the bone marrow of *Thbs1^+/+^* or *Thbs1^-/-^* mice and stimulated them with LPS (100 ng/mL) and IFNγ (20 ng/mL) for 4 hours. The stimulated *Thbs1^+/+^* or *Thbs1^-/-^* Mφs were co-cultured, in a Transwell chamber, with a mouse aortic SMC line (MOVAS) for 24 hours. qPCR analyses of MOVAS cells showed that SMCs co-cultured with *Thbs1^-/-^* Mφs had lower abundance of *Thbs1* mRNA compared to SMCs co-cultured with *Thbs1^+/+^* Mφs (Figure S5C).

### SMC phenotypes were altered by myeloid *Thbs1* gene deficiency

Although DEG analysis showed increased abundance of genes related to SMC contraction, immunostaining showed a significant reduction in SMC-specific MYH11 in the AngII-stimulated *Thbs1*^ΔMφ^ aortic tissues (Figure 5A). To understand this apparent discrepancy, we further examined the six SMC subclusters (Figure 5B). Based on the content of enriched genes associated with clusters, we classified SMC-2 as matrix-producing, SMC-3 proliferative, SMC-4 contractile, SMC-5 inflammatory or Mφ-like, and SMC-6 hemoglobin positive. SMC-1, the biggest subcluster, were deemed to be intermediate SMCs based on the abundance of both contractile and matrix genes (Figure 5C, Figure S6). Compared to wildtype, *Thbs1*^ΔMφ^ aortas contained increased SMC-1 relative to SMC-2, 3, and 4 (Figure 5D). Since SMC-1 expressed moderate levels of *Myh11* compared to SMC-4, the expansion of SMC-1 and shrinkage of SMC-4 may explain the reduction of MYH11 abundance detected by immunostaining.

**Figure 5.**
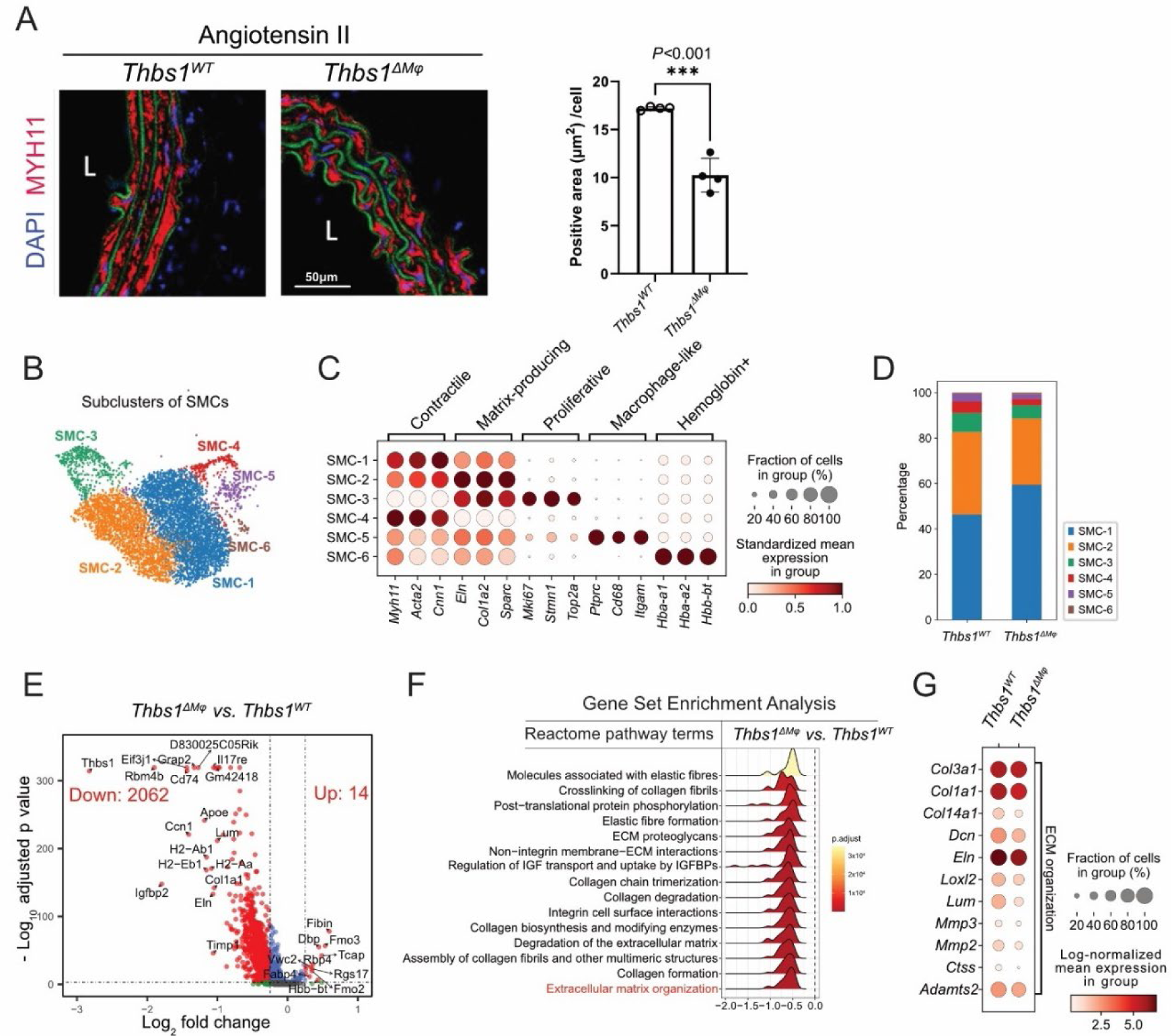
Myeloid *Thbs1* deficiency altered smooth muscle cell (SMC) phenotypes. **(A)** Immunostaining and quantification of aortic cross-sections after 7 days of Ang II infusion using anti-MYH11. Green: elastin autofluorescence. “L” indicates the aortic lumen. **(B)** UMAP plot of sub-populations in SMCs of *Thbs1^WT^* and *Thbs1^ΔMφ^* data combined. **(C)** Expression of enriched genes in each sub-population. **(D)** Percentage of sub-populations in *Thbs1^WT^* and *Thbs1^ΔMφ^*. **(E)** Volcano plot of differentially expressed genes (DEGs) in total SMCs (*Thbs1^ΔMφ^* versus *Thbs1^WT^*). **(F)** Gene Set Enrichment Analysis (GSEA) of the altered gene sets in *Thbs1^ΔMφ^* versus *Thbs1^WT^* group. **(G)** Expression of genes in ECM organization pathway in *Thbs1^WT^* and *Thbs1^ΔMφ^*.

When examined the transcriptomic changes in SMCs, we observed there were a much greater number of downregulated, than upregulated, genes (2062 genes downregulated vs 14 genes upregulated) (Figure 5E). Pathway analysis showed that multiple downregulated gene sets are related to ECM synthesis and organization such as *Col1a1, Eln, and Thbs1* (Figure 5F&G).

### Alteration of fibroblasts and myofibroblasts caused by myeloid *Thbs1* gene deficiency

Fibroblasts are important for repairing ECM in addition to SMCs. We identified four fibroblast subclusters and three myofibroblast subclusters (Figure 6A). Based on their gene expression profiles, we inferred them as stem cells antigen-1 positive (SCA1+) progenitor fibroblasts (Fib-1), ECM producing fibroblasts (Fib-2), ECM cross-linking fibroblasts (Fib-3), proliferative fibroblasts (Fib-4), proliferative myofibroblasts (Myofib-1), ECM producing myofibroblasts (Myofib-2), and EC-like myofibroblasts (Myofib-3) (Figure 6B). While the total fibroblast percentage decreased in *Thbs1*^ΔMφ^ aortas (Figure 3D), noticeable shifts took place in the subcluster composition: *Thbs1*^ΔMφ^ aorta contained lower percentage of Fib-1 and slightly higher Fib-4. In contrast, all three myofibroblast subclusters expanded in *Thbs1*^ΔMφ^, while the total myofibroblast percentages were similar between *Thbs1*^ΔMφ^ and wildtype aortas (Figure 6C). Similar to what was noted in SMCs, DEG analysis further showed that 2253 genes were downregulated, and only 47 genes were upregulated in fibroblasts and myofibroblasts in *Thbs1*^ΔMφ^ compared to the wildtype aortas (Figure 6D). The downregulated genes were primary genes that encode ECM proteins such as *Col1a1*, *Col3a1*, and upregulated genes contained many contractile genes such as *Cnn1*, *Myh11*, *Acta2*, and *Tagln* (Figure 6D).

**Figure 6.**
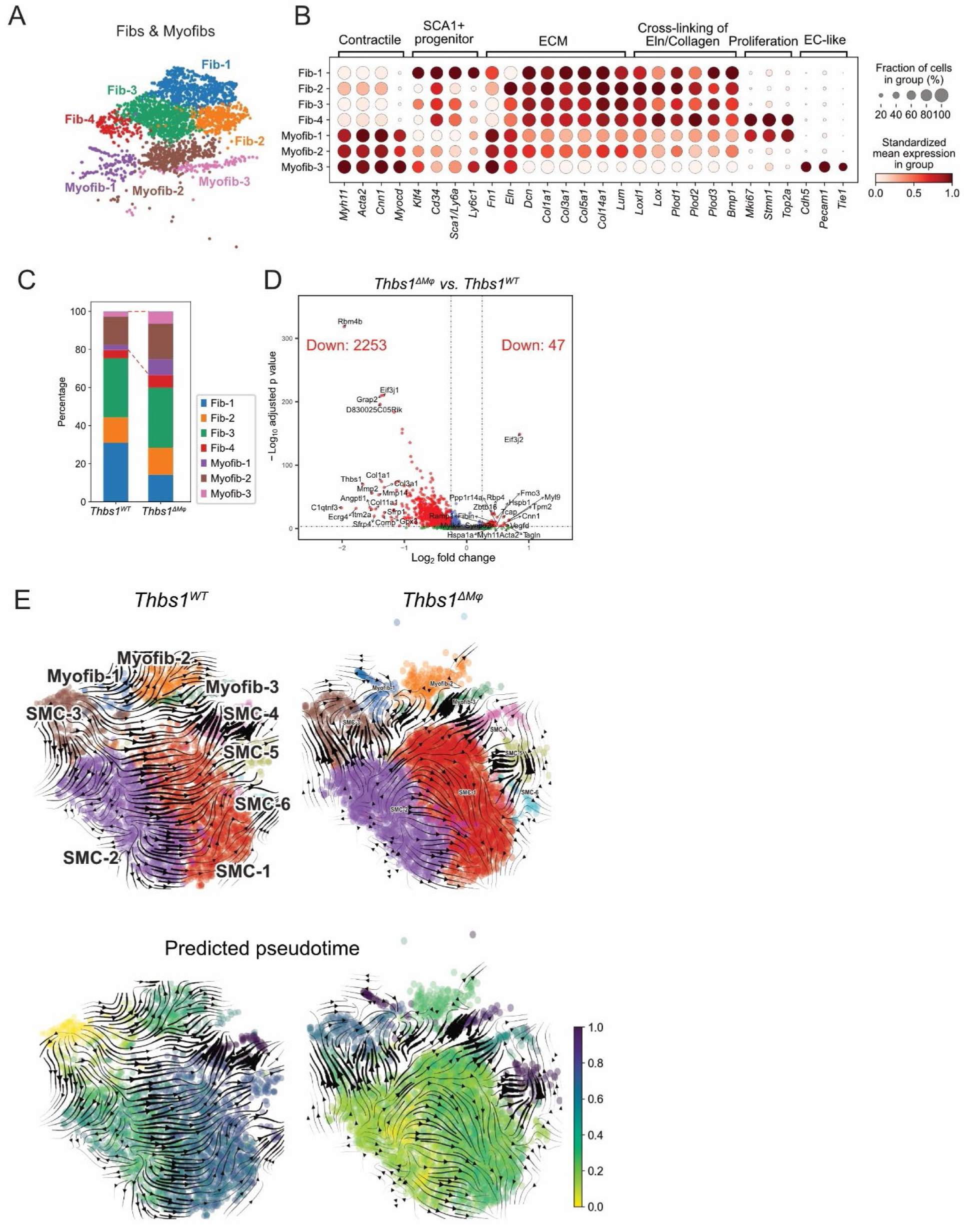
Myeloid *Thbs1* deficiency altered fibroblasts and myofibroblasts. **(A)** UMAP plot of sub-populations in fibroblasts and myofibroblasts of *Thbs1^WT^* and *Thbs1^ΔMφ^* data combined. **(B)** Expression of enriched genes in each sub-population. **(C)** Percentage of sub-populations in *Thbs1^WT^* and *Thbs1^ΔMφ^*. **(D)** Volcano plot of differentially expressed genes (DEGs) in total fibroblasts and myofibroblasts (*Thbs1^ΔMφ^* versus *Thbs1^WT^*). **(E)** RNA velocity analysis of smooth muscle cells (SMCs), fibroblasts, and myofibroblasts in *Thbs1^WT^* and *Thbs1^ΔMφ^*.

To understand how ECM producing cells became less abundant in the mutant aortic wall, we further analyzed the phenotypic transition among SMCs and myofibroblasts by inferring the RNA velocity. As shown in Figure 6E, there was less phenotypic transition among subclusters in *Thbs1*^ΔMφ^ compared to wildtype aortas. In wildtype, proliferative SMC (SMC-3) was the starting point in the RNA velocity-predicted pseudotime, and changing towards matrix-producing SMC and myofibroblasts (SMC-2 and Myofib-2), and finally contractile SMC (SMC-4). In *Thbs1*^ΔMφ^, matrix-producing SMC (SMC-2) transitioned into proliferative SMC (SMC-3), and finally Mφ-like SMC (SMC-5) and EC-like myofibroblasts (Myofib-3).

### Cell-Cell communications were reduced in *Thbs1*^ΔMφ^ mice

To understand the molecular mechanism responsible for the defected ECM production in *Thbs1*^ΔMφ^ aorta, we performed CellChat^24^ analysis on the scRNA-seq data. Comparing to the AngII-incubated wildtype aortas, AngII-incubated *Thbs1*^ΔMφ^ aortas had lower ligand-receptor interactions and interaction strength (Figure 7A&B). The detected ligand-receptor pairs among the 8 major cell types fell into 30 signaling pathways (Figure 7C), 6 of which were altered by myeloid-specific *Thbs1* deficiency. Major histocompatibility class I (MHC−I), transforming growth factor-β (TGFβ), protein S (PROS), and non-canonical Wnt (ncWNT) pathways were downregulated, whereas signal regulatory protein (SIRP) and C-C motif chemokine ligand (CCL) pathways upregulated in the mutants (Figure 7C). TGFβ signaling was one of the most decreased pathways in *Thbs1*^ΔMφ^ compared to the wildtype (Figure 7C and Figure S7A). Closer examination of the TGFβ pathway revealed that the three ligands were expressed by different cell types: *Tgfb1* by dendritic cells and Mφs, *Tgfb2* by SMCs and myofibroblasts, and *Tgfb3* by SMCs, fibroblasts and myofibroblasts (Figure 7D). Myeloid-specific *Thbs1* deficiency profoundly reduced the abundance of *Tgfb3* (Figure 7D). Consequently, Tgfb3 – (Acvr1+Tgfbr1), which regulates fibroblast-to-SMC, myofibroblast-to-SMCs, and SMC-to-SMC communications in the wildtype, became undetectable in *Thbs1*^ΔMφ^ (Figure 7E).

**Figure 7.**
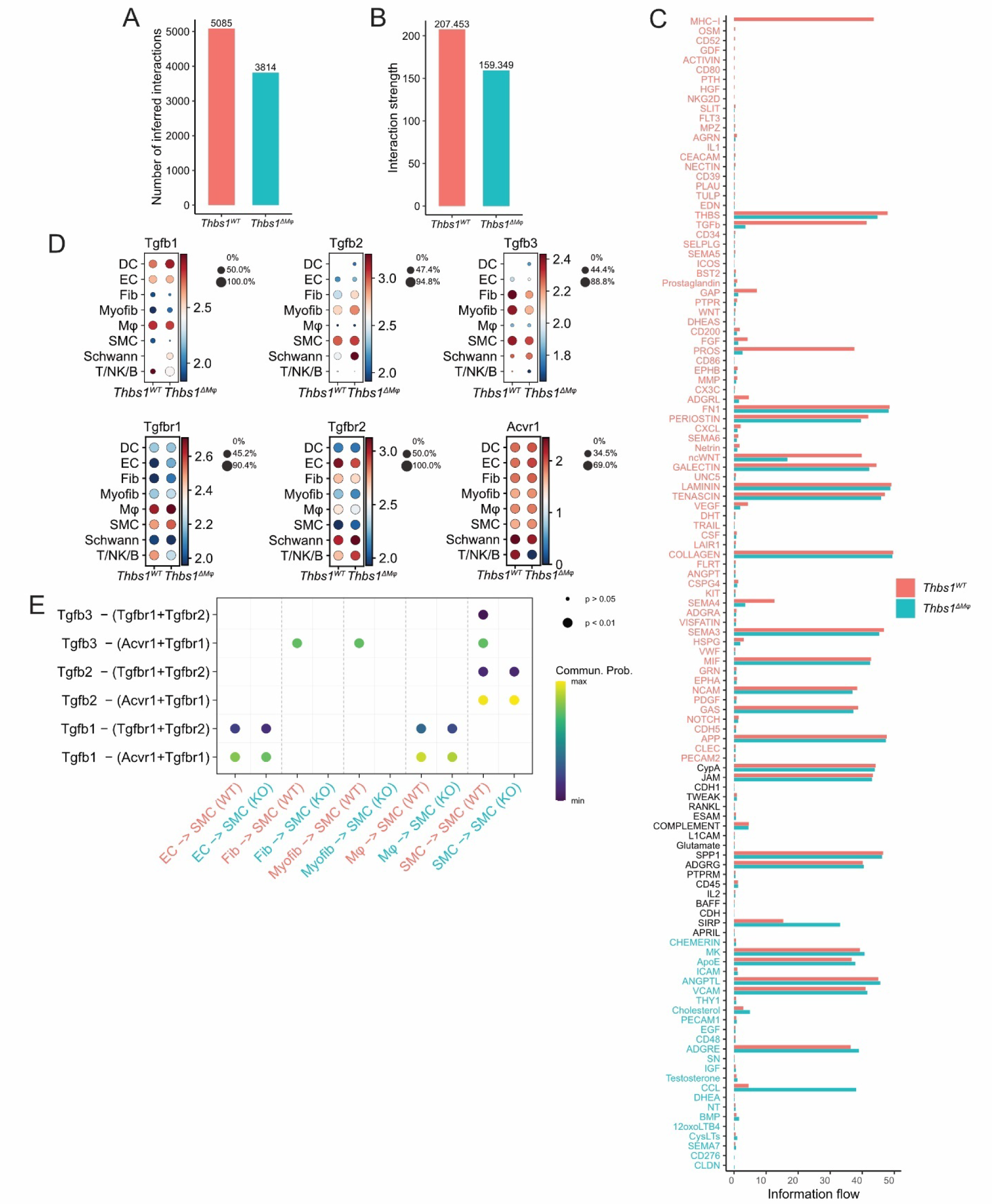
Cell-cell communications in rupture-prone aortic tissues. **(A&B)** Number of predicted ligand-receptor pairs and total interaction possibility in *Thbs1^WT^* and *Thbs1^ΔMφ^* groups. **(C)** Information flow of each signaling pathway in *Thbs1^WT^* and *Thbs1^ΔMφ^* groups. **(D)** Expression of ligands and receptors of TGFβ signaling in each cell type in *Thbs1^WT^* and *Thbs1^ΔMφ^* groups. **(E)** Communication probability of each ligand-receptor pairs that contributed to TGFβ signaling sent from endothelial cells (EC), fibroblasts (Fib), myofibroblasts (Myofib), macrophages (Mφ) and smooth muscle cells (SMCs) to SMCs in *Thbs1^WT^* (WT) and *Thbs1^ΔMφ^* (KO) groups.

MHC−I signaling was only detected in the wildtype mice, and it communicated among ECs, Mφs, and dendritic cells (DC) (Figure S7B). PROS signaling driven by EC, fibroblasts, and myofibroblasts, and ncWNT signaling dominant by fibroblasts and myofibroblasts were also decreased in *Thbs1*^ΔMφ^ (Figure S7C&D). In contrast, SIRP and CCL pathways were enhanced in *Thbs1*^ΔMφ^, specifically in DCs (Figure S8).

### Reduced *THBS1* expression at dissection sites in human AAA

To enhance translational insights, we performed Visium spatial transcriptomic analysis on aortic tissues obtained from AAA patients who underwent emergency rupture repair (referred to as the emergency group, n=4) or elective aneurysm repair (referred to as the elective group, n=4). Specimens from both patient groups exhibited severe abnormalities in H&E stains, such as atherosclerotic plaques, calcifications, inflammatory cell accumulation, and arterial disorganization (Figure 8A). Unbiased analysis via Chrysalis revealed multiple “compartments” in each sample, in which a compartment refers to a distinct region or area within tissue that exhibits unique gene expression patterns. Figure 8A and Figure S9-11 plotted all compartments in each sample and the top 20 enriched genes in each compartment. There was considerable heterogeneity among samples within each group. Due to the great heterogeneity, data from each sample had to be analyzed individually, and could not be combined and analyzed together as in the scRNA-seq experiments. As such, no significant difference in gene expression was found between the aortic tissues of patients undergoing emergency rupture repair and those undergoing elective AAA repair. Figure 8A showcased analysis of an aortic section from emergency group, displaying 8 compartments with distinct gene expression patterns.

**Figure 8.**
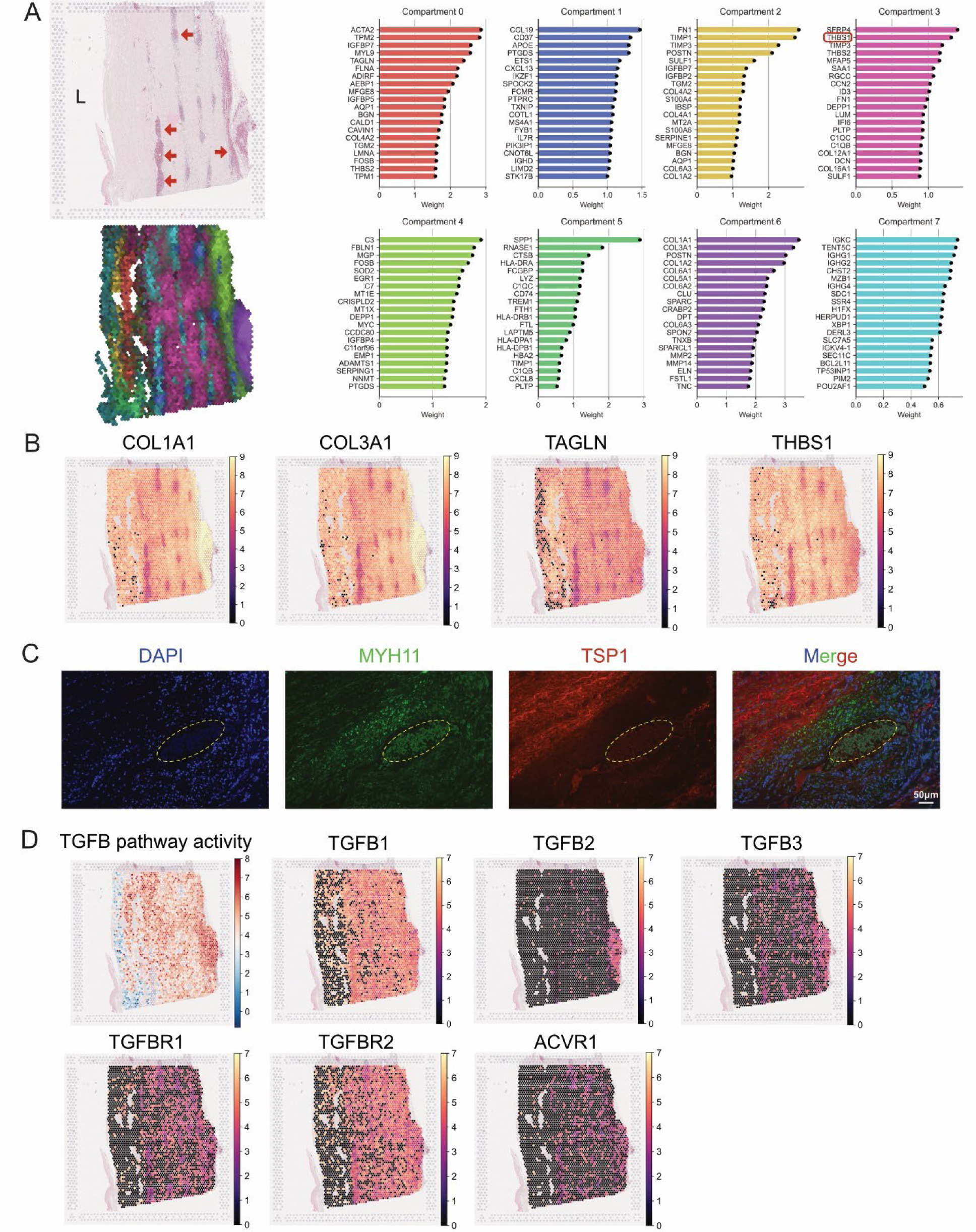
Spatial transcriptomic analysis of human abdominal aortic aneurysm (AAA) tissues. **(A)** Representative H&E staining and tissue compartment analysis with the top 20 enriched genes in each compartment. *THBS1* was highlighted by red rectangles. Red arrow indicates dissection sites (presence of red blood cells). “L” indicates the aortic lumen. **(B)** Expression and distribution of *COL1A1*, *COL3A1*, *TAGLN*, and *THBS1* in representative AAA tissue. **(C)** Immunostaining of human AAA cross-sections with anti-MYH11 and anti-TSP1. The yellow dashed ellipse indicates the dissection site. **(D)** TGFβ (TGFB) pathway activity and expression of ligands and receptors of TGFβ signaling in AAA tissue.

All samples contained dissection sites/compartments marked by the presence of red blood cells in H&E stains and enrichment of immunoglobulin genes (IGHGs). By comparing gene abundance between dissection and non-dissection sites within a single specimen, we found dissection sites had lower *THBS1* and matrix gene expression such as *COL1A1* and *COL3A1*, and *TAGLN*, compared to non-dissection sites (Figure 8B). Immunostaining on human AAA tissues confirmed that dissection sites had lower TSP1 abundance (Figure 8C). Further analysis on TGFβ signaling also showed that TGFB3 and ACVR1 abundances were higher at dissection sites (Figure 8D).

## Discussion

In this manuscript, we reveal that myeloid-specific deficiency of TSP1 significantly exacerbates aortic rupture, primarily through mechanisms involving broad suppression of ECM proteins. By employing murine models with cell-specific deletion of TSP1, we showed that myeloid deficiency of *Thbs1* increased aortic rupture rates by more than twofold. The pro-rupture function of TSP1 appeared to be restricted to myeloid cells most likely Mφs because endothelial-specific *Thbs1* knockouts did not show this effect. ScRNA-seq and histological analyses identified a unique cellular and molecular signature associated with rupture-prone aortas, characterized by diminished inflammation and reduced ECM production. Furthermore, spatial transcriptomic analysis of human AAA tissues showed a correlation between low *THBS1* expression and aortic dissection. Collectively, these results suggest that TSP1 from Mφs plays an essential role in aortic tissue repair by promoting ECM synthesis within the aneurysmal aorta, thereby affords intrinsic resistance to aneurysm rupture.

Mφ infiltration is an important pathophysiological characteristic of aortic aneurysm. The diminished presence of Mφs in aortas from AngII-infused *Thbs1*^ΔMɸ^ mice was consistent with our previously reported function of TSP1 in Mφ migration and tissue invasion^11,12^. Unexpectedly, reductions in leukocyte accumulation coincided with lethal aortic rupture of *Thbs1*^ΔMφ^ mice, which contradicts the conventional belief that Mφs contribute to aneurysm progression by releasing ECM degrading proteases and pro-apoptotic proinflammatory cytokines^28,29^. The role of Mφs in aortic deterioration is supported by experimental evidence showing that inhibition of inflammatory infiltration, such as gene deletion of CCR2, renders aneurysm resistance and protect ECM and SMCs against destruction^30,31^. However, the low inflammatory state in AngII-infused *Thbs1*^ΔMφ^ mice is clearly noticeable and demonstrated by two independent methods, scRNA-seq and by immunostaining. The low abundance of F4/80 positive cells in *Thbs1^Δ^*^Mφ^ aortic tissues persisted from an early (3 days) to late (28 days) intervals during the course of AngII infusion. The ultrasound measurement of aortic diameter at baseline, day 3 and day 7 showed mice of both phenotypes dilated their aorta to a similar degree before rupture incidence took place. However, the aortic diameters of mice survived to day 28 were smaller in the *Thbs1^Δ^*^Mφ^ group. These multifaceted phenotypes resulting from myeloid-specific *Thbs1* gene deficiency suggest that the role of Mφs in aneurysm progression and rupture might be more complex than previously thought. While Mφ-mediated inflammation is a critical pathological driver of aortic dilation, Mφs, likely a subpopulation of Mφs, might be necessary for the intrinsic vascular repair mechanisms. Unfortunately, the *Lyz2-cre* causes gene deletion in all myeloid linages including the pro-healing Mφs. The lack of both pro-aneurysm and pro-healing Mφs may explain the prone-rupture and small aortic dilation phenotypes of *Thbs1^Δ^*^Mφ^ mice.

Since ∼60% of *Thbs1^Δ^*^Mφ^ aortae ruptured between day 7 and day 14 during AngII-infusion, we suggest that the transcriptomic and histological analyses on day 7 tissues provided clues to potential mechanisms that exasperate aortic rupture in *Thbs1^Δ^*^Mφ^ mice. Indeed, scRNA-seq data showed that loss of TSP1 in myeloid cells had a broad effect on the gene expression of ECM proteins as well as the ECM-producing cells. Across several cell populations within the aortic wall of *Thbs1^Δ^*^Mφ^ mice, gene sets related to ECM proteins, or their regulators were markedly reduced compared to the wildtype mice. Both *Col1a1* and *Col1a2* are among the 10 most downregulated genes. Although it is technically challenging to definitively prove the causal effect of ECM *Thbs1^Δ^*^Mφ^ rupture, plentiful evidence in the literature supports the key role of ECM in maintaining vascular tissue integrity and in aneurysm pathophysiology^32^. Inherited vascular connective tissue disorders predispose patients to early onset of aortic aneurysm, dissection, and rupture as seen in Marfan Syndrome and Loeys-Dietz Syndrome^33,34^. In mice, genetic manipulation of genes coding for ECM proteins or proteins regulates ECM synthesis and assembly causes or exacerbates aortic rupture and/or dissection^35–42^. SMCs and fibroblasts are the major source of ECM proteins, however, not all SMCs and fibroblasts are equal in producing ECM proteins. Among the 6 subclusters of SMCs and 7 subclusters of fibroblasts, SMC-2 and Fib-1 are mostly enriched with transcripts of ECM proteins or related proteins including elastin, Col1a2, and PAI-1 in SMC-2 and Col1a1 and Col3a1 in Fib-1. The shrinking of SMC-2 and Fib-1 cell populations in *Thbs1^Δ^*^Mφ^ is a plausible explanation why there is a broad reduction in the expression of ECM related genes. The molecular signature of SMC-2 cluster is similar to a cell cluster called fibromyocytes that were identified in human ascending aortic tissues by Li and colleagues: high expression of contractile genes as well as ECM genes^43^. Since the cell death pathway was not significantly altered by *Thbs1^Δ^*^Mφ^, apoptosis or necrosis are the unlike cause of reduction in SMC-2 and Fib-1 cells. Consistent with the notion that SMCs undergo phenotypic change during aneurysm development^44–47^, our RNA velocity analysis suggests proliferative SMCs (SMC-3) were the origin of other SMCs in wildtype mice, whereas in *Thbs1^Δ^*^Mφ^, matrix-producing SMCs (SMC-2) might assume the phenotype of other SMC subclusters. This phenotypic switch between SMCs subclusters could be responsible for the redistribution of SMC subtypes and subsequent reduction in ECM-making SMCs.

The analysis of cell-cell communication by CellChat casts light on the molecular mechanism that may regulate the phenotypic shift of SMCs and fibroblasts. Among the differentiated expressed signaling pathways, the TGFβ pathway was a focus because of its known functions in regulating SMC phenotypes, ECM synthesis and aneurysm pathophysiology^48–50^. Consistent with this literature, we found by the CellChat analysis that *Thbs1* mutant aortas displayed a profound loss of Tgfb3–(Acvr1 and Tgfbr1) mediated communication from fibroblasts/myofibroblasts to SMCs or SMCs to SMCs, which is likely because of diminished expression the Tgfb3 gene. However, additional work is needed to determine whether Tgfb3 is responsible for maintaining ECM producing cells in the aortic wall.

TSP1 may contribute to aneurysm pathophysiology through several mechanisms in addition to regulating Mφ infiltration. TSP1 contributes to TGFβ activation through its interaction with latent TGFβ complex^51^. Disrupting the interaction between TSP1 and latency-associated peptide with an antagonistic peptide, Krishna and colleagues was able to produce a larger aortic aneurysm with enhanced inflammation, reduced TGFβ signaling, and more severe ECM degradation in AngII-treated *Apoe^-/-^* mice^52^. Another potential mechanism for TSP1 to promote aneurysm is through its functions in endothelial cells. While aneurysm research has been largely focused on inflammatory cells and SMCs, the contribution of endothelial dysfunction to aneurysm is increasingly appreciated^8,53^. EC-specific deletion of *Thbs1* moderately reduced the aneurysm size. Ultrasound imaging showed a better recovery of aortic compliance following aneurysm reduction in the *Thbs1^Δ^*^EC^ relative to the wildtype mice. In the current study, we chose to dive deep into the mechanisms underlying the prone-to-rupture property of *Thbs1^Δ^*^Mφ^ in order to address the knowledge gap regarding aneurysm rupture. Future effort will be devoted to detailed analysis of the aneurysm protective phenotype of *Thbs1^Δ^*^EC^ mice.

Our observation of region-specific TSP1 expression in human AAA tissues may provide an explanation for the controversial literature regarding TSP1 abundance in aortic aneurysmal diseases. By comparing TSP1 immunostaining in human and mouse aortic tissues with or without aneurysms, our group as well as Yamashiro and colleagues found that TSP1 abundance was upregulated in AAA^11,12^ and thoracic aortic aneurysm^54^. However, Krishna and collegues showed an reduced presence of TSP1 in AAA body tissues compared with the relatively normal AAA neck^55^. The current data from Visium analysis of human AAA tissues is consistent with findings by Krishna et al, highlighting the region dependent and perhaps cell dependent expression pattern of TSP1. Therefore, the contributions of TSP1 to the pathophysiology of aortic aneurysm need to be examined at regional even cellular level.

There are several limitations in this study. As mentioned earlier in the discussion, *Lyz2-cre* leads to deletion of *Thbs1* in monocyte/Mφs, granulocytes and some CD11c+ dendritic cells^56^. Although we demonstrated previously that TSP1 did not directly impact polarization of Mφs, both recruited bone marrow-derived and resident Mφs expressed TSP1. It is currently unclear whether resident Mφs, as well as TSP1 expressed by these cells, are pro- or anti-aneurysm. Another limitation is related to the AngII model, which produces many pathological features of human AAAs. The AngII model also produces aortic dissection, a risk factor of aneurysm rupture^44^. We were unable to accurately detect aortic dissection due to an instrumental limitation.

In conclusion, this study identifies myeloid TSP1 as a key player in preventing AAA rupture. Mice lacking TSP1 specifically in myeloid cells exhibited a significantly increased rate of rupture, likely through impairing extracellular matrix production. Targeting mechanisms that regulate myeloid TSP1 expression could hold promise for novel therapeutic strategies to prevent AAA rupture.

## Acknowledgement

We thank Dr. Nader Sheibani for generously providing the *Thbs1*^ΔEC^ and *Thbs1* flox mice, and Jooyong Kim for technical assistance. Images of immunofluorescent and RNA *in situ* hybridization were captured at UW Madison Optical Imaging Core. Single-cell RNA sequencing was performed at UW Biotechnology Center DNA Sequencing Facility. Spatial transcriptomics was performed at UW Translational Research Initiatives in Pathology (TRIP) Lab and UW Biotechnology Center Gene Expression Center. Histology of mouse tissues was performed at the UW Histology Resource Core. Ultrasound imaging and analysis was conducted at UW Small Animal Imaging & Radiotherapy Facility.

## Supplemental Material

Figure S1-S11.

