## Supplemental Material for "Myeloid-Specific Thrombospondin-1 Deficiency Exacerbates Aortic Rupture via Broad Suppression of Extracellular Matrix Proteins"

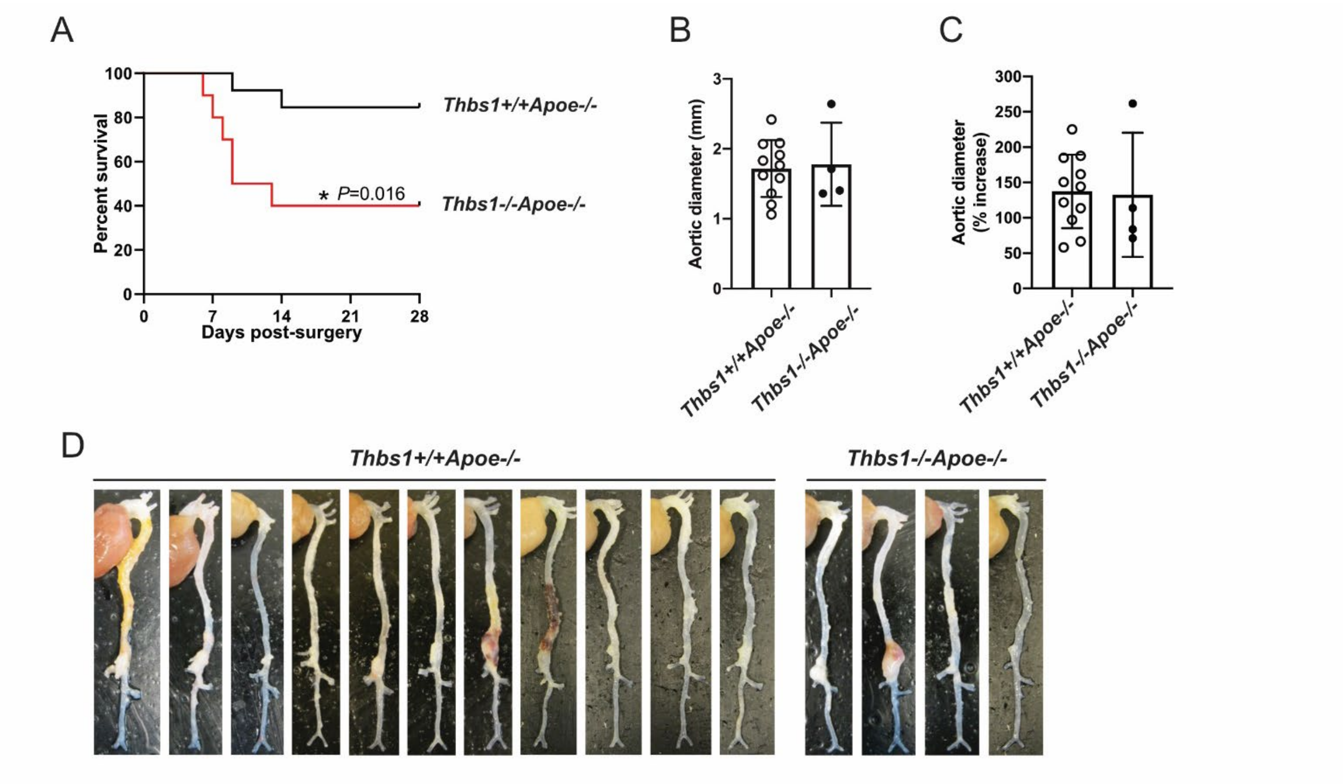


**Figure S1. Global TSP1 deficiency increased aneurysm rupture in angiotensin II-infused mice. (A)** Survival curve of *Thbs1*$^{+/+}$/*Apoe*$^{-/-}$ and *Thbs1*$^{-/-}$/*Apoe*$^{-/-}$ mice during 28 days of angiotensin II (Ang II) infusion (1000 ng/kg/min). **(B&C)** Maximum external aortic diameter at the suprarenal abdominal aorta in survived mice (B), and the percentage increase compared to the maximum external aortic diameter of the infrarenal region (C). **(D)** Images of the aortas taken during euthanization on Day 28 of Ang II infusion. A total of 13 *Thbs1*$^{+/+}$/*Apoe*$^{-/-}$ and 11 *Thbs1*$^{-/-}$/*Apoe*$^{-/-}$ were subjected to Ang II infusion. Log-rank test was performed in (A). *$P$<0.05. Data in (B) and (C) were presented as mean±SD; two-tailed Student t test was performed.

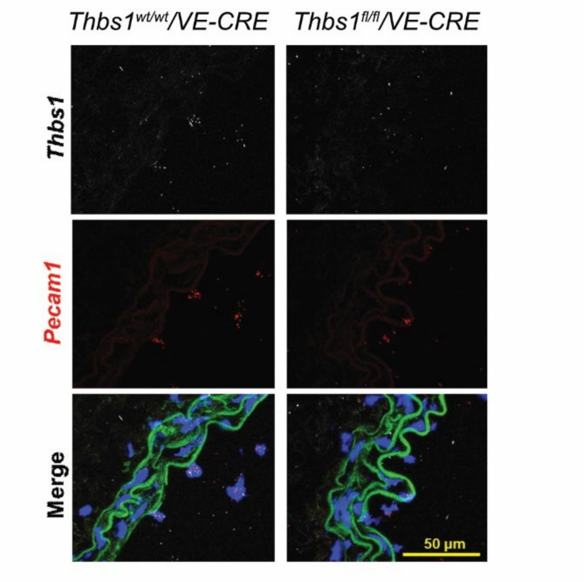


**Figure S2. Validation of endothelial-specific *Thbs1* deficient mice.** RNA fluorescent *in situ* hybridization (RNA-FISH) of *Thbs1* and *Pecam1* on aortic cross-sections of *Thbs1*$^{wt/wt}$*/VE-Cadherin Cre* (*Thbs1*$^{wt/wt}$*/VE-CRE*) and *Thbs1*$^{flox/flox}$*/VE-Cadherin Cre* (*Thbs1*$^{fl/fl}$*/VE-CRE*) mice. Representative images from 3 independent samples were shown. Green: elastin autofluorescence.

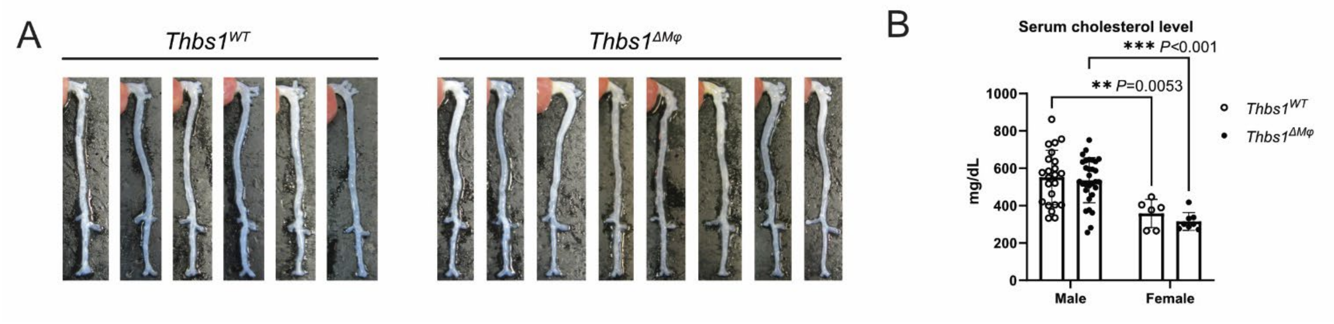


**Figure S3. Female *Thbs1*$^{WT}$ and *Thbs1*$^{\Delta M\varphi}$ mice did not develop aneurysm in AAV. PCSK9D377Y-angiotensin II (Ang II) model. (A)** Images of the aortas in both groups. **(B)** Serum cholesterol levels of male and female mice who developed hypercholesterolemia in response to AAV.PCSK9D377Y (serum cholesterol > 250 mg/dL), 14 days after AAV injection. N=6 for *Thbs1*$^{WT}$ and n=8 for *Thbs1*$^{\Delta M\varphi}$ were subjected to Ang II infusion. Data in (B) were presented as mean±SD. Two-way ANOVA followed by Tukey's multiple comparisons was performed. **$P$<0.01, ***$P$<0.001.

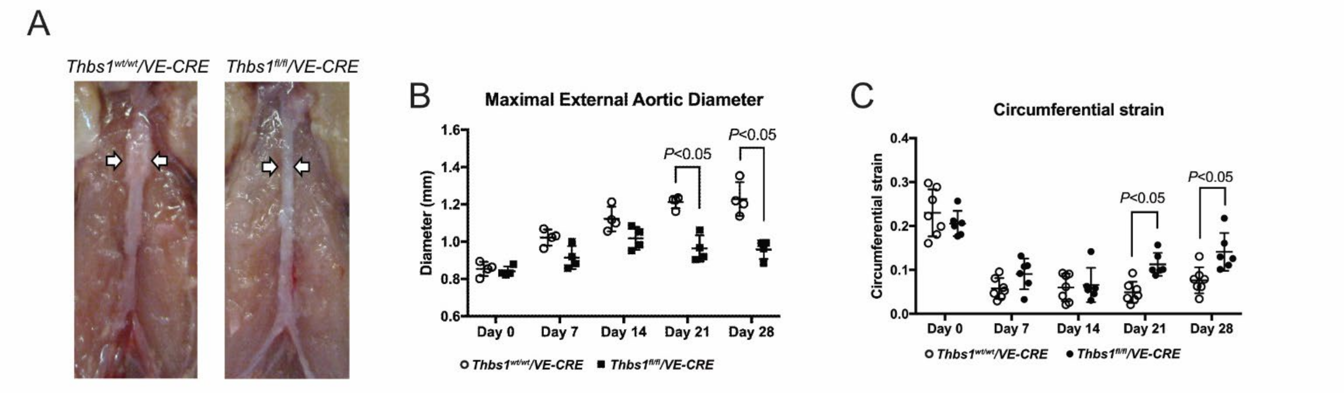


**Figure S4. Endothelial-specific *Thbs1* deficiency reduced abdominal aortic aneurysm (AAA) in $CaCl_2$ model. (A)** Representative images of the injured infrarenal abdominal aortas in both groups. **(B)** Maximum external aortic diameter at the infrarenal abdominal aorta measured by ultrasound imaging. **(C)** Circumferential strain of the infrarenal abdominal aorta calculated analyzed by ultrasound imaging. Data in (B) and (C) were presented as mean±SD. Two-way ANOVA followed by Tukey's multiple comparisons was performed.

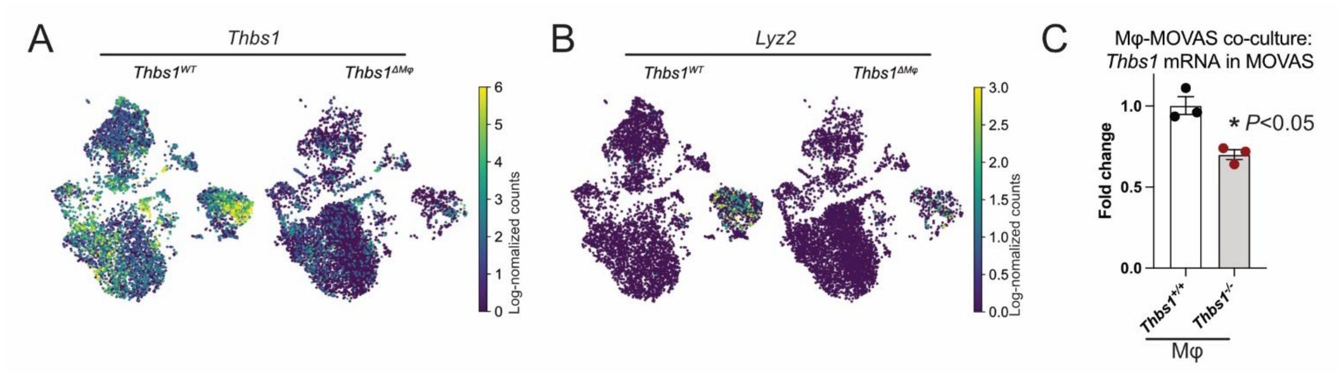


**Figure S5. *Thbs1* deficiency in macrophages decreased *Thbs1* expression in smooth muscle cells. (A&B)** *Thbs1* (A) and *Lyz2* (B) expression and distribution in *Thbs1*$^{WT}$ and *Thbs1*$^{\Delta M\varphi}$ aortic tissues. **(C)** Bone marrow-derived macrophages from *Thbs1*$^{+/+}$ and *Thbs1*$^{-/-}$ mice were stimulated with 100ng/mL LPS and 20ng/mL IFNγ for 4 hours, then co-cultured with mouse aortic smooth muscle cell line (MOVAS) in transwell setting for 24 hours. Levels of *Thbs1* mRNA in MOVAS were determined by real-time PCR (polymerase chain reaction). Data were presented as mean±SD of at least 3 independent experiments. Two-tailed Student t test was performed. *$P<0.05$.

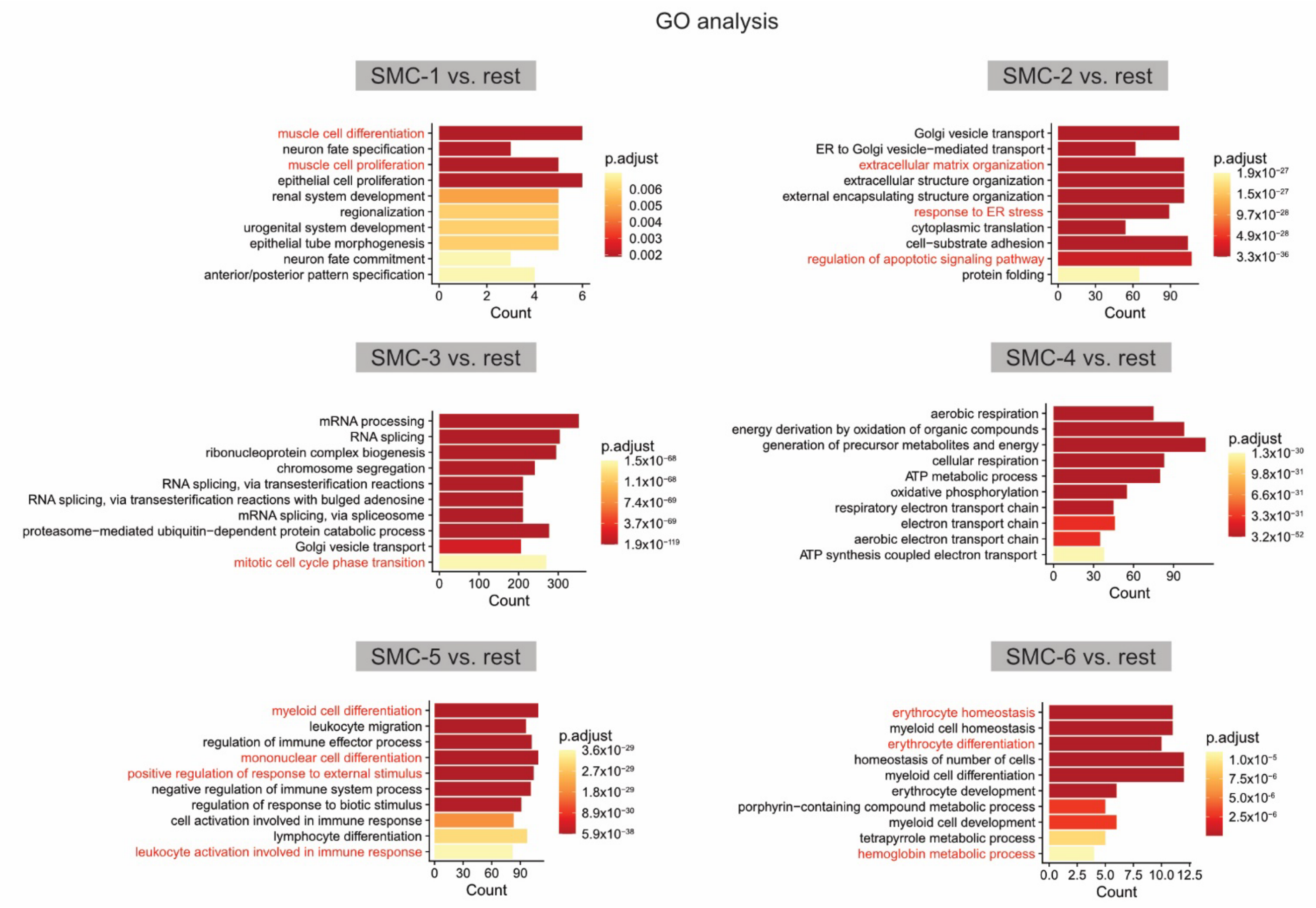


**Figure S6.** Gene Ontology (GO) analysis of each smooth muscle cell sub-population in comparison to the other 5 sub-populations.

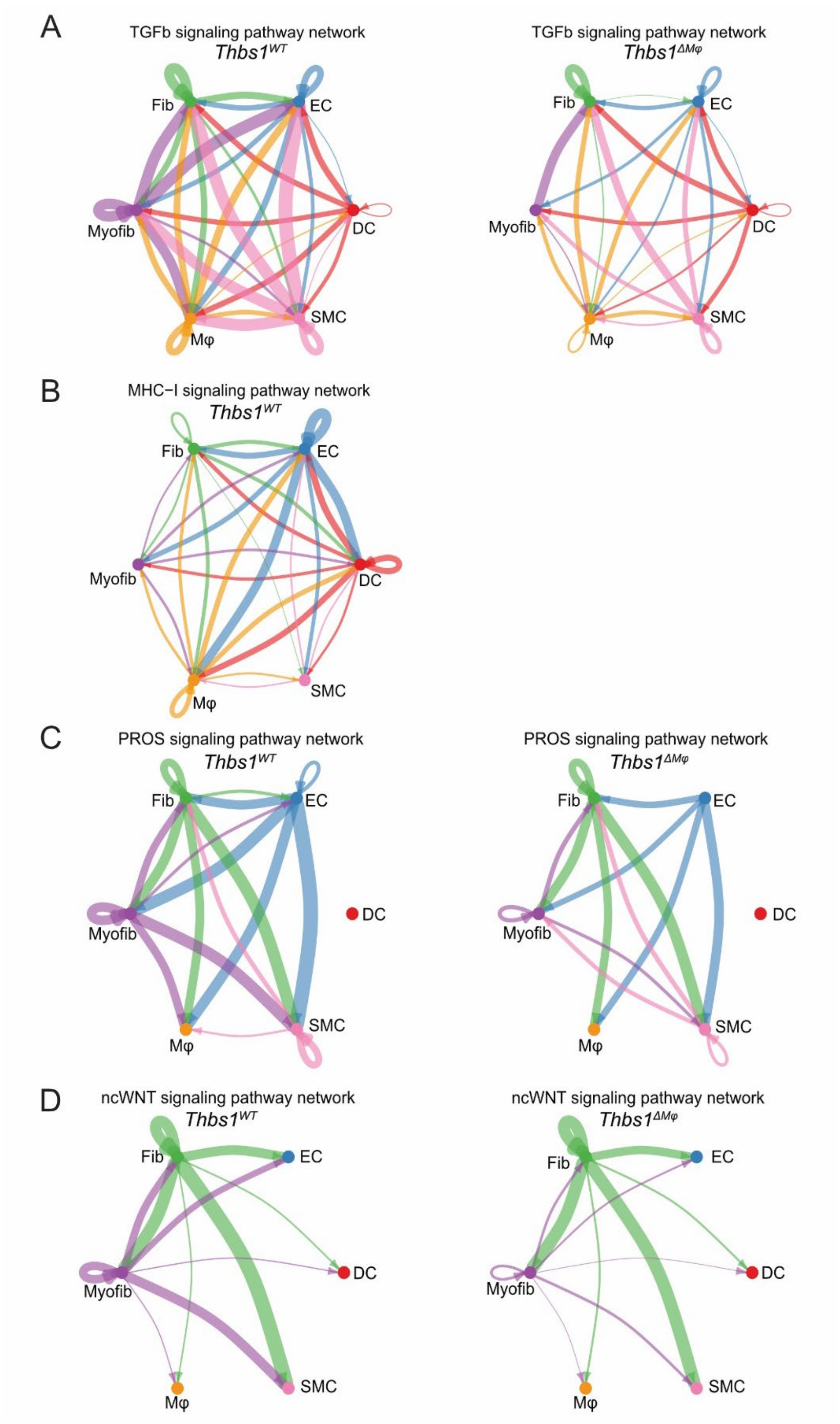


**Figure S7.** Circle plot of TGFβ (TGFb) **(A)**, MHC-I **(B)**, PROS **(C)**, ncWNT **(D)** signaling network in *Thbs1*$^{WT}$ and *Thbs1*$^{\Delta M\varphi}$ groups.

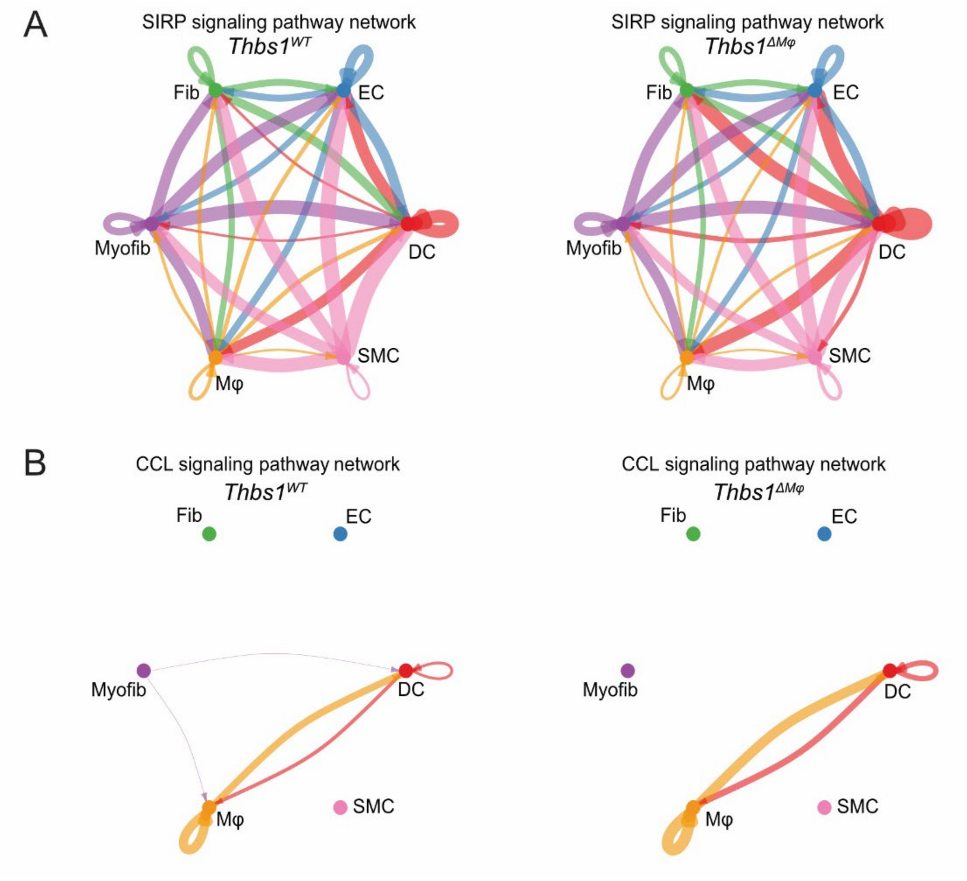


**Figure S8.** Circle plot of SIRP **(A)** and CCL **(B)** signaling network in *Thbs1*$^{WT}$ and *Thbs1*$^{\Delta M\varphi}$ groups.

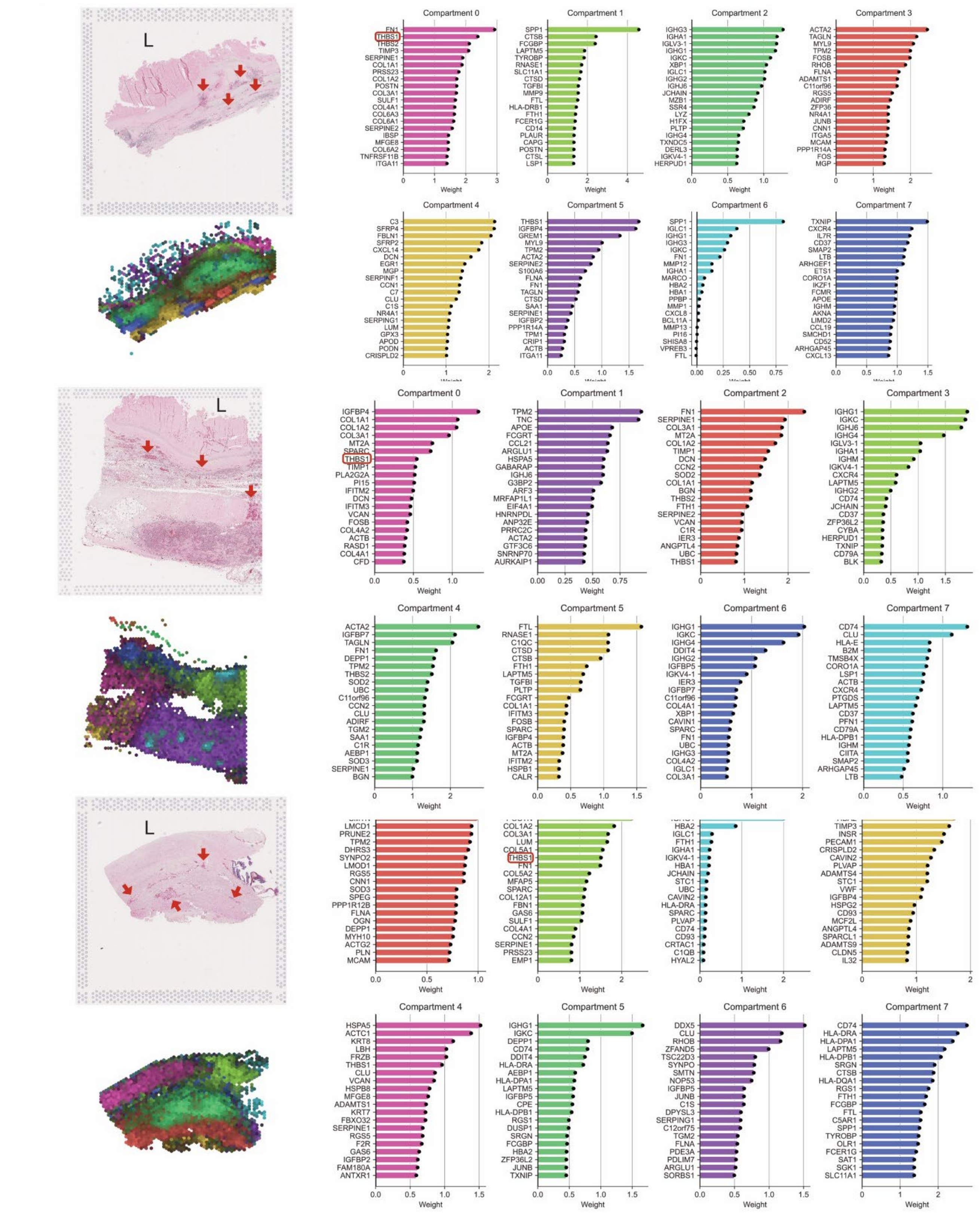


**Figure S9.** H&E staining and tissue compartment analysis of human abdominal aortic aneurysm (AAA) tissues. *THBS1* was highlighted by red rectangles. "L" indicates the aortic lumen.

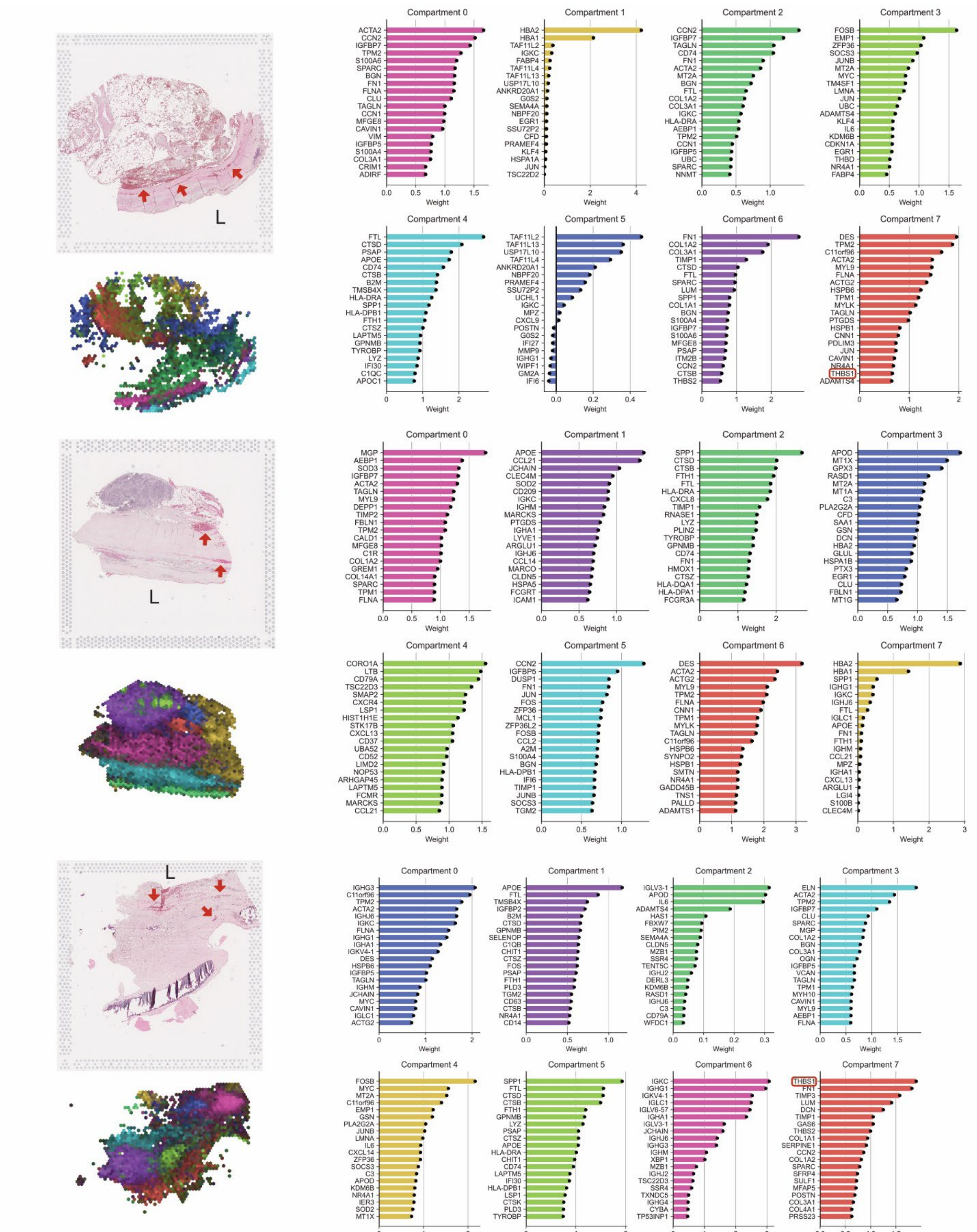


**Figure S10.** H&E staining and tissue compartment analysis of human abdominal aortic aneurysm (AAA) tissues. *THBS1* was highlighted by red rectangles. “L” indicates the aortic lumen.

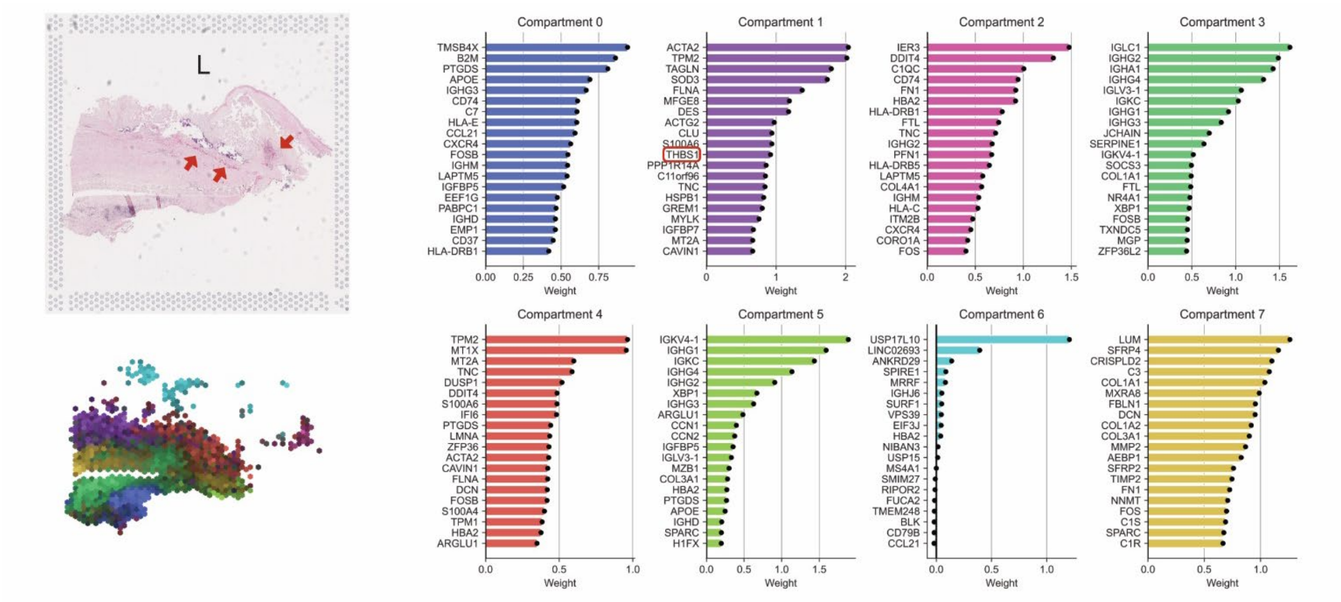


**Figure S11.** H&E staining and tissue compartment analysis of human abdominal aortic aneurysm (AAA) tissues. *THBS1* was highlighted by red rectangles. "L" indicates the aortic lumen.
